# Ammonium regulates the development of pine roots through hormonal crosstalk and differential expression of transcription factors in the apex

**DOI:** 10.1101/2021.07.15.452498

**Authors:** Francisco Ortigosa, César Lobato-Fernández, Hitomi Shikano, Concepción Ávila, Shu Taira, Francisco M. Cánovas, Rafael A. Cañas

## Abstract

Ammonium is a prominent source of inorganic nitrogen for plant nutrition, but excessive amounts can be toxic for many species. However, most conifers are tolerant to ammonium, a relevant physiological feature of this ancient evolutionary lineage. For a better understanding of the molecular basis of this trait, ammonium-induced changes in the transcriptome of maritime pine (*Pinus pinaster* Ait.) root apex have been determined by laser capture microdissection and RNA sequencing. Ammonium promoted changes in the transcriptional profiles of multiple transcription factors, such as *SHORT-ROOT*, and phytohormone-related transcripts, such as *ACO*, involved in the development of the root meristem. Nano-PALDI-MSI and transcriptomic analyses showed that the distributions of IAA and CKs were altered in the root apex in response to ammonium nutrition. Taken together, the data suggest that this early response is involved in the increased lateral root branching and principal root growth, which characterize the long-term response to ammonium supply in pine. All these results suggest that ammonium induces changes in the root system architecture through the IAA-CK-ET phytohormone crosstalk and transcriptional regulation.

## INTRODUCTION

Nitrogen (N) is a vitally important nutrient for all living organisms because it is a constituent of essential biomolecules such as nucleic acids, proteins, amino acids, porphyrins and hormones, among others (Miller & Cramer, 2005). This nutrient is indispensable for the proper growth and development of plants, and it can be assimilated from different kinds of sources, including organic (peptides, amino acids, and urea) and inorganic (nitrate and ammonium) forms (Näsholm et al., 1998, Hachiya & Sakakibara, 2017). Together with nitrate (NO_3_^-^), ammonium (NH_4_^+^) is one of the main forms of inorganic N available for plants, and the relative proportions of these elements in soil can vary depending on biological and climate conditions (Bijlsma et al., 2000).

Experimental evidences strongly suggest that NH_4_^+^ is perceived and recognized by plant cells as a signal that promotes physiological and morphological changes in plants (Liu and von Wirén, 2017). However, NH_4_^+^ at millimolar levels usually causes toxicity in most plants. The effects caused by an excessive NH_4_^+^ availability include decreased plant growth, leaf chlorosis, decreased root/shoot ratios, decreased root gravitropism and altered root system architecture (RSA) (Esteban et al., 2016). Regarding the changes in RSA, the inhibition of root elongation, enhancement of lateral root (LR) branching and impaired root gravitropism are commonly observed (Y. Liu & von Wirén, 2017).

Root elongation involves three interconnected biological processes: cell division, cell expansion and cell differentiation (Youssef et al., 2018). Both cell division and cell expansion are affected by NH_4_^+^ through two independent determinants: i) a decreased capacity for protein N-glycosylation, and ii) an increased production of reactive oxygen species (ROS) (Jia & von Wirén, 2020). In *Arabidopsis,* root growth inhibition in response to NH_4_^+^ represses root cell production by decreasing the meristem size and the number of dividing cells without altering the cell division rate (Y. Liu et al., 2013). Furthermore, NH_4_^+^ also decreases the number of root cap cells (Y. Liu et al., 2013). In this sense, several transcription factors (TFs) have been described to play key roles in root cap development, such as BEARSKIN (BRN) and SOMBRERO (SMB) (Bennett et al., 2010, 2014; Kamiya et al., 2016). However, the transcriptional pathway regulating NH_4_^+^ root cap development has not yet been deciphered.

In addition, auxins (IAAs) play a prominent role in cell division and cell expansion (Wang & Ruan, 2013). IAAs can be synthesized in the root meristem and transported toward upstream adjacent regions through specific transporters, such as AUXIN RESPONSE 1 (AUX1) and PINFORMED 1-7 (PIN1-7) (Grunewald & Friml, 2010). IAAs induce the expression of PLETHORA 1-4 (PLT1-4) TFs in the root meristem, which are responsible for cell proliferation maintenance (Galinha et al., 2007). Other TFs have been described to play key roles in the stem cell niche and the quiescent center (QC) maintenance to produce all tissues required to form a mature root, including the TFs SHORT-ROOT (SHR) and SCARECROW (SCR) (Sablowski, 2011). *SHR* expression is localized in a zone of the stele that constitutes the central part of the root and stem (Miyashima et al., 2011; Kim et al., 2020). When *SHR* transcripts are translated, SHR proteins move into adjacent cells (pericycle cells, endodermis, QC and phloem pole) to activate the expression of SCR (Helariutta et al., 2000; Nakajima et al., 2000; Sena et al., 2004; Cui et al., 2007). SHR also plays key roles in phloem development, controlling the asymmetric cell division process for sieve element development by the regulation of NARS1 and SND2 NAC-type TFs (Kim et al., 2020). This fact is relevant because these cell types constitute the phloem (Greb, 2020), which is the main means of IAA transport (Chapman & Estelle, 2009). However, Y. Liu et al. (2013) showed that NH_4_^+^ does not affect the IAA content at QC, where the maximum level of this phytohormone in roots is localized (Zhou et al., 2010).

As it was mentioned above, LR branching is enhanced by NH_4_^+^ supply (Araya et al., 2016). When the quadruple ammonium transporter (AMT) knockout lines, that exhibit a severe reduction in LR branching, were complemented with *AtAMT1.1* or *AtAMT1.3*, only *AtAMT1.3*-complemented plants were able to restore the LR branching phenotype induced by NH_4_^+^ (Lima et al., 2010), suggesting that the NH_4_^+^-induced LR branching signaling events are dependent on AtAMT1.3 (Lima et al., 2010). Recently, it has been demonstrated that under NH_4_^+^ supply, RSA changes mediated by IAAs transport are highly linked to root acidification (Jia et al., 2020; Meier et al., 2020), which seems to be the molecular basis for the AMT transporter effect on LR branching since NH_4_^+^ uptake is accompanied by pH imbalance. In addition, IAAs have been suggested to be involved in this signaling event due to the repression of PIN2 under a supply of NH_4_^+^ (Y. Liu et al., 2013; Zou et al., 2013), which is provoked by the PIN2 hyperphosphorylation (Ötvös et al., 2021). Regarding root agravitropic response, NH_4_^+^ promotes the downregulation of AUX1 and PIN2, which are two pivotal IAA transporters and are subject to the antagonistic action between PIN2 and ARG1. ARG1 is involved in the transduction of the root gravity signal and required for normal AUX1 expression and basipetal IAA transport in root apices, promoting an asymmetric auxin flow (Y. Liu et al., 2013; Zou et al., 2013). All these evidences highlight the importance of IAA in orchestrating the root system configuration in response to this nutritional stimulus. However, the transcriptional regulatory mechanisms that control this response remain unclear.

Together with IAAs, cytokinins (CKs) have been previously established to be important in root plant growth and vascular development (Kamada-Nobusada et al., 2013; Miyashima et al., 2019; Mao et al., 2020). CK biosynthesis and activity in plants are closely related to N availability (Takei et al., 2004; Kamada-Nobusada et al., 2013). In the roots of rice, NH_4_^+^ nutrition leads to the accumulation of different CKs and CK-derived compounds (Kamada-Nobusada et al., 2013). During active growth of *Arabidopsis*, the content of CKs first increased in the vascular system, thus reflecting CKs transport from roots to shoots (Shtratnikova et al., 2015). Recently, it has been reported that CK signaling in the early protophloem-sieve-element cell files of *Arabidopsis* root procambial tissue promotes the expression of several DOF TFs (Miyashima et al., 2019). Together with the IAA-responsive dependent HD-ZIP III proteins, these TFs compose a transcriptional network that integrates spatial information of the hormonal domains and miRNA gradients, which are essential for root vascular development (Miyashima et al., 2019).

Previous works in *Arabidopsis*, based on root protoplast generation and cell-sorting using FACS coupled to expression studies, revealed cell-specific responses to different processes such as N nutrition (Gifford et al., 2008; Walker et al., 2017) or plant immunity (Rich-Griffin et al 2020). Regarding N nutrition, co-expression network approaches revealed that within the root cortex N controlled a wide range of processes including phytohormone responses, while in pericycle regulatory networks were related to formation of new organs, organ structure development, and establishment of cell localization (Walker et al., 2017) showing that phytohormones are candidates signaling for cell-specific responses to N (Gifford et al., 2008).

Conifers are an ancient group of gymnosperms that cover vast regions in the Northern Hemisphere, and they have exceptional ecologic and socioeconomic importance (Farjon et al., 2018). Since conifers represent a differentiated evolutionary lineage of plants, they exhibit substantial differences in N metabolism with respect to angiosperms such as *Arabidopsis*. For instance, conifers lack glutamine biosynthesis in the plastid of photosynthetic cells (Cánovas et al., 2007), and most species prefer NH_4_^+^ over NO_3_^-^ as the main source of inorganic N (Kronzucker et al., 1997; Hawkins & Robbins, 2010). This is the case for maritime pine (*Pinus pinaster* Aiton) (Warren & Adams, 2002; Ortigosa et al., 2020), a southwestern Mediterranean conifer employed as a model in studies of N nutrition and metabolism and for which a large body of genomics resources are available (Cañas et al., 2017; Ortigosa et al., 2020; 2021). Previous transcriptomic studies of pine seedling tissues provided an overview of the gene expression distribution in different pine tissues but also it was possible to highlight the relationships between gene expression patterns and function in a tissue-dependent manner (Cañas et al., 2014; 2017).

Maritime pine seedlings under NH_4_^+^ supply accumulate more biomass than those fed with NO_3_^-^ what is related with a higher uptake of NH_4_^+^ than NO_3_^-^ (Ortigosa et al., 2020). In maritime pine roots, a long-term supply of NH_4_^+^ induced the expression of transcripts related to defense, such as *antimicrobial peptide 1* (*PpAMP1*) (Canales et al., 2010), which is not found in dicots and can regulate the NH_4_^+^ uptake (Canales et al., 2011). There also was a close relationship between NH_4_ ^+^-responsive genes and genes involved in amino acid metabolism, particularly those involved in asparagine biosynthesis and utilization (Canales et al., 2010). Additionally, NH_4_^+^ promotes changes in the epitranscriptome that mainly regulate the translational response and growth, including the repression of 1-aminocyclopropane-1-carboxylic acid (ACC) oxidase (ACO), the last enzyme in the ethylene (ET) biosynthesis pathway (Ortigosa et al., 2021). Considering this background, the main goal of the present work was to decipher the molecular mechanisms underlying the early response to NH_4_^+^ in the root apex and its relationship with root development in maritime pine.

## MATERIALS AND METHODS

### Plant material

Maritime pine seeds (*P. pinaster* Aiton) from “Sierra Segura y Alcaraz” (Albacete, Spain) were provided by the *Área de Recursos Genéticos Forestales* of the Spanish *Ministerio de Agricultura, Pesca y Alimentación*. Seed germination was carried out following the protocol described elsewhere (Cañas et al., 2006). Seedlings were grown in vermiculite in plant growth chambers (Aralab Fitoclima 1200, Rio de Mouro, Portugal) under 16Lh light photoperiod, a luminal intensity of 125LμmolLm Ls, a constant temperature of 23L°C, 50% relative humidity and watered twice a week with distilled water. One-month-old maritime pine seedlings were used for the experiments. Pine seedlings were randomly subdivided into three different groups, relocated into forestall seedbeds and irrigated with 80 mL of a solution that contains macro- and micronutrients without any N source (1.16 mM KCl; 0.63 mM KH_2_PO_4_; 0.35 mM MgSO_4_·7H_2_O; 0.17 mM CaCl_2_·H_2_O; 80 μM EDTA-FeSO_4_; 25.9 μM H_3_BO_3_; 10.2 μM MnCl_2_·4H_2_O; 1.3 μM ZnSO_4_·7H_2_O; 0.7 μM CuSO_4_·5H_2_O; 0.1 μM Na_2_MoO_4_·2H_2_O). After three days of acclimation, the control group was irrigated with 80 mL of water and the experimental group was supplied with 80 mL of 3 mM NH_4_Cl. Root samples were collected 24 h after the treatment and immediately frozen in liquid N (Figure 1).

**Figure 1.**
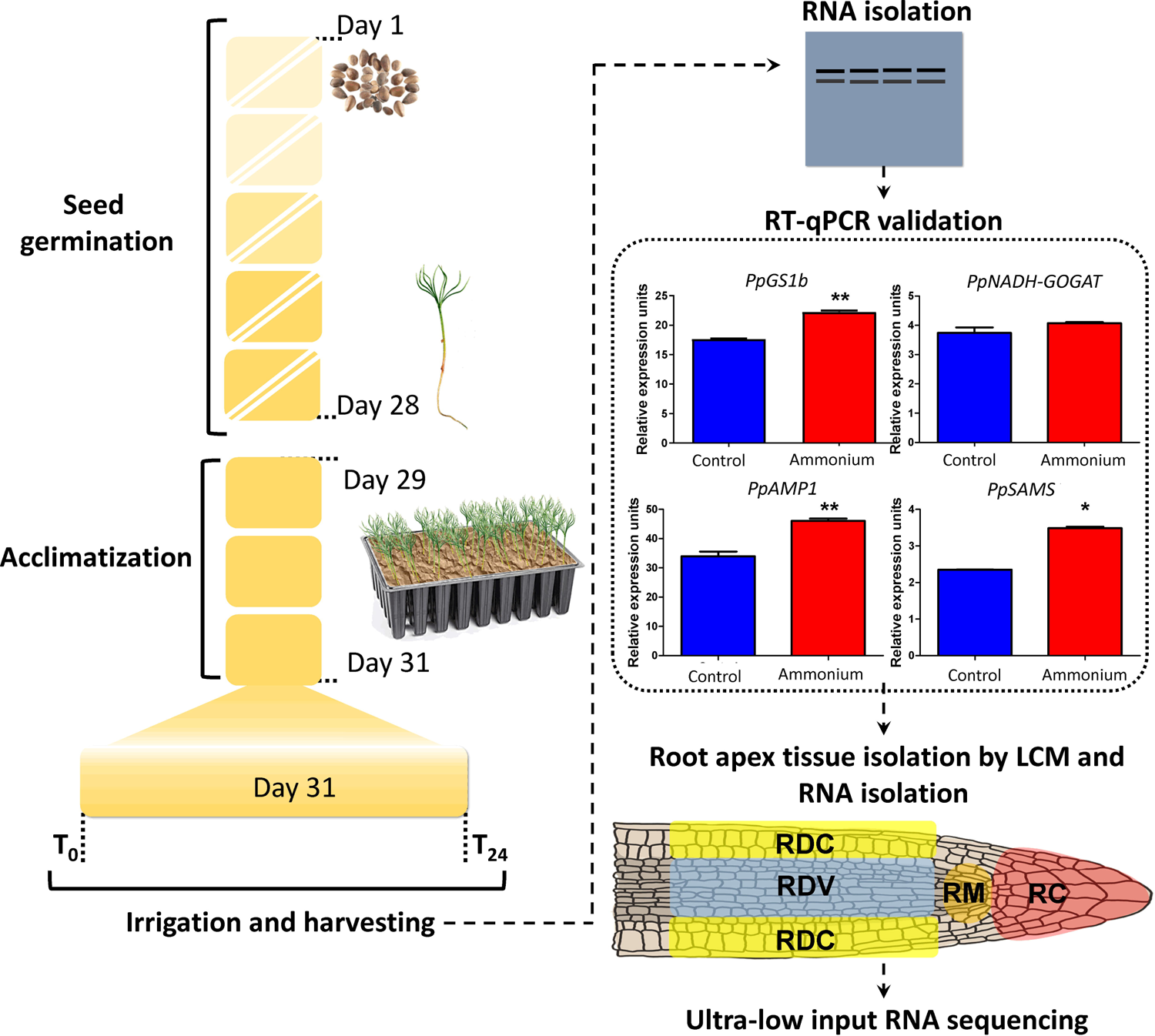
Experimental design. Timeline from germination to harvest. RT-qPCR data for the validation of the experimental design are included. *Glutamine synthetase 1b* (*PpGS1b*); *NADH-dependent glutamate synthase* (*PpNADH-GOGAT*); *antimicrobial peptide 1* (*PpAMP1*); *S-adenosyl methionine synthase* (*PpSAMS*) Significant differences were determined with a *t*-test (* *P*-value < 0.05; ** *P*-value < 0.01). Error bars show SE with n = 3. Isolated regions from the root apex: cap (RC), meristem (RM), developing cortex (RDC) and developing vessels (RDV).

Ten seedlings were collected and pooled per each sample for RT-qPCR validation. The same experiment was carried out three independent times. The screening condition and adequate development of each experiment was verified through the gene expression analysis by RT-qPCR of control genes following previous results (Ortigosa et al., 2021) (Figure 1). To perform laser capture microdissection (LCM), seedling’s root apexes were cut, and tissues (5-6 mm) were imbibed in a specimen holder with Tissue-Tek optimal cutting temperature embedding medium (Sakura Finetek, Alphen aan den Rijn, The Netherlands) and immediately frozen in liquid N for cryostat sectioning. Frozen samples were stored at −80°C until use.

### Root phenotyping in response to nitrogen supply

For the root phenotypes under different nitrogen sources (NH_4_ and NO_3_), the pine seedlings were cultivated in a growth chamber with a 16/8 h light/dark photoperiod, light intensity of 125 μmolL^−2^ s^−1^, constant temperature of 23°C, and 50% relative humidity (Aralab Fitoclima 1200, Rio de Mouro, Portugal). One-month-old seedlings (n = 30) were randomly transplanted into seedbeds for each experimental condition: distilled water, 3 mM KNO_3_ and 3 mM NH_4_Cl. Each experimental condition group was irrigated twice per week with 50 mL of the corresponding N solution or 50 mL of distilled water for 74 days. After treatment, seedling roots were dissected and whole roots, primary root (PR) and LRs were weighted. The PR length and number of LRs was manually assessed (Figure S1).

### Whole root RNA isolation

Total RNA from roots of maritime pine seedlings was isolated following the protocol described by Liao et al. (2004) and modified by Canales et al. (2012). RNA concentration and purity were determined spectrophotometrically using a Nanodrop ND-1000 (Thermo Scientific, Waltham, MA, USA). Purity of the preparation was assessed by determining the 260/280 and 260/230 ratios. The integrity was assessed by electrophoresis.

### Laser capture microdissection, RNA isolation and low-input RNA-seq

LCM procedure was carried through as previously described (Cañas et al., 2014). Full step protocol is described in Methods S1. Four different tissue areas were isolated by microdissection corresponding to the root cap (RC), meristem (RM), developing cortex (RDC) and developing vessels (RDV) areas (Video S1).

All RNA extractions from the microdissection procedure were carried out using manufacturer’s instruction protocol (non-LCM) for the RNAqueous-Micro RNA Isolation Kit (Ambion, Austin, TX, USA). RNA quality, DNA contamination and first quantification were performed via RNA Pico Assay for the 2100 Bioanalyzer (Agilent, Santa Clara, CA, USA). Quantification was verified via a Qubit RNA BR (Broad-Range) Assay Kit (Invitrogen, Paisley, UK). RNA samples with RNA integrity number (RIN) higher than 7 were used for subsequent RNA sequencing, mRNA amplification and cDNA synthesis 1 ng of RNA for each sample.

The low input RNA-seq was carried out by Novogen (Hong Kong). The cDNA synthesis and amplification, and the library preparation was made with the SMART-Seq™ v4 Ultra™ Low Input RNA Kit for Sequencing (Takara, Mountain View, CA, USA) following the manufacturer’s instructions. RNA sequencing was made in a NovaSeq 6000 sequencer according to the manufacturer’s instructions for paired-end reads (Illumina, San Diego, CA, USA). The 24 samples were sequenced producing paired-end reads of 150 bp length. The sequencing output is shown in the Table S1.

**Table 1.** Most representative DE transcripts coding for TFs and hormone related proteins.

| TF in<br>maritime pine | Gene ID | Tissue | Transcript in<br>*A.thaliana* | Gene ID | Process involved | References |
| --- | --- | --- | --- | --- | --- | --- |
| *PpDOF11* | *pp_58878* | RC | *HCA2/*<br>*DOF5.6* | *AT5G62940* | Root tip vascular proliferation: radial growth of protophloem sieve elements | Guo et al., 2009<br>Miyashima et al., 2019 |
| *PpSHR* | *pp_246622* | RC | *SHR* | *AT4G37650* | - Quiescent center cells specification<br>- Asymmetric cell division related to rootradial pattern formation<br>- Phloem development | Fukaki et al., 1998<br>Helariutta et al., 2000<br>Cui et al., 2007<br>Kim et al., 2020 |
| *PpGATA20-*<br>*like* | *pp_204819* | RC | *GATA20* | *AT2G18380* | Root tip vascular development | Smit et al., 2020 |
| *PpSGR/IDD*<br>*16* | *pp_174625* | RC | *IDD14* | *AT1G68130* | - Lateral organ morphogenesis and gravitropic responses<br>- Establishment of auxin gradients<br>(auxin biosynthesis and transport) | Cui et al., 2013 |
| *PpPHR1-*<br>*like* | *pp_64350* | RD<br>C | *PHR1* | *AT4G28610* | Phosphate starvation | Rubio et al., 2001 |
| *PpOFP17* | *pp_80824* | RD<br>C | *OFP17* | *AT2G30395* | Plant growth and development | Wang et al., 2011 |
| *PpBLISTER-*<br>*like* | *pp_170664* | RD<br>C | *BLISTER* | *AT3G23980* | Plant vegetative growth | Schatlowskiet al., 2010<br>Hong et al., 2019 |
| *PpDOF2* | *pp_119525* | RD<br>C | *PEAR1/*<br>*DOF2.4* | *AT2G37590* | Root tip vascular proliferation: radial growth of protophloem sieve elements | Miyashima et al., 2019 |
| *PpDOF12* | *pp_85381* | RM | *CDF3* | *AT3G47500* | Involved in abiotic stress tolerance and nitrogen response | Corrales et al., 2017<br>Domínguez-Figueroa et al., 2020 |
| *PpDOF1* | *pp_233816* | RM | *OPB3* | *AT3G55370* | Regulator of phyB and cry1 signaling pathways | Ward et al., 2005 |
| *PpHAT-like* | *pp_141831* | RM | *HAT1* | *AT4G17460* | - Inflorescence meristem regulation, floral meristem determination, gynoecium medial domain development<br>- Repressor of BR responsive genes and ABA biosynthesis | Zúñiga-Mayo et al., 2012<br>Zhang et al., 2014<br>Tan et al., 2018 |
| *PpINCURV*<br>*ATA-like* | *pp_85012* | RM | *ATBH-15* | *AT1G52150* | Root meristem patterning and putatively in the regulation of procambial and vascular tissue formation or maintenance. | Ochando et al., 2006 |
| *PpNAC31* | *pp_168080* | RM | *SMB* | *AT1G79580* | Cellular maturation of root cap regulation | Willemsen et al., 2008<br>Bennett et al., 2010<br>Fendrych et al., 2014 |
| *PpNAC38* | *pp_72282* | RM | *BRN1/BRN2* | *AT1G33280* | Cellular maturation of root cap regulation | Bennett et al., 2010 |
| *PpVAN3/FKD2* | *pp_60103* | RC<br>and<br>RM | *VAN3/*<br>*FORKED2* | *AT5G13300* | Plant vascular tissue formation by PIN transporters localization | Steynen & Schultz, 2003<br>Naramoto et al., 2009<br>Hou et al., 2010 |
| *Cytokinin riboside*<br>*5'-monophosphate*<br>*phosphoribohydrolase* | *pp_222714* | RC | *LOG3* | *AT2G37210* | Cytokinin activation | Kuroha et al., 2009 |
| *Cycloeucaleanol cycloisomerase* | *pp_189071* | RC | *CPII* | *AT5G50375* | PIN2 endocytosis | Men et al., 2008 |
| *Sulfotransferase* | *pp_208269* | RDC | *SULT20B1* | *AT3G45070* | Flavonoids sulfation | Hashiguchi et al., 2013 |
| *PpABA2* | *pp_198945* | RDC and RM | *ABA2* | *AT1G52340* | Biosynthesis of abscisic acid | González-Guzmán et al., 2002 |
| *PpCYP90B1* | *pp_247190* | RDC | *CYP90B1* | *AT3G50660* | Biosynthesis of brassinosteroids | Fujita et al., 2006 |
| *PpTIR1* | *pp_238715* | RDC | *TIR1* | *AT3G62980* | Auxin signalling pathway | Dharmasiri et al., 2005<br>Salehin et al., 2015 |
| *PpCYP87A3* | *pp_81114* | RDC | *CYP87A2* | *AT1G12740* | Putatively involved in hormone biosynthesis and/or auxin signaling | Chaban et al., 2003 Bark et al., 2011 |
| *GASA gibberellin regulated cysteine richprotein* | *pp_111379* | RDC | *GASA10* | *AT5G59845* | Putatively involved in response to phytohormones | Zhang & Wang, 2017 |
| *HMG-CoA reductase* | *pp_144669* | RDC | *HMGR* | *AT2G17370* | Regulatory enzyme of the cytokinin and brassinosteroid biosynthesis pathways | Antolín-Llovera et al., 2011 |
| *PpABA2* | *pp_209084* | RDC | *ABA2* | *AT1G52340* | Biosynthesis of abscisic acid | González-Guzmán et al., 2002 |
| *Cytokinin dehydrogenase* | *pp_215376* | RDC | *CKX1* | *AT2G41510* | Catalyzes the oxidation of cytokinins | Werner et al., 2003 |
| *PpACO* | *pp_202153* | RM | *ACO* | *AT2G19590* | Ethylene biosynthesis | Houben et al., 2019 |
| *Flavonoid 3'-monoxygenase* | *pp_203885* | RM | *F3'H* | *AT5G07990* | Involved in flavonoids biosynthesis pathway from naringenin to eriodictyol and dihydrokaempferol to dihydroquercetin | Schoenbohm et al., 2000 |
| *EXORDIUM* | *pp_129078* | RM | *EXO* | *AT4G08950* | Putatively involved in brassinosteroids response in roots | Coll-Garcia et al., 2004 |

The raw reads were trimmed (quality and contamination) using SeqTrimBB software (https://github.com/rafnunser/seqtrimbb). Only the pairs in which both reads passed the quality test were further analyzed (Q>20). Trimmed reads are shown in the Table S1. These reads were assembled using Trinity 2.11 (Haas et al., 2013). Contigs lower than 400 pb were eliminated, for the rest of contigs the redundancy was reduced using CD-HIT-EST software (Fu et al., 2012). This transcriptome shotgun assembly project has been deposited at DDBJ/EMBL/GenBank under the accession GJFX00000000. The version described in this paper is the first version, GJFX01000000. The final transcriptome was used as the reference for the read mapping that was performed with BWA using the MEM option (Li and Durbin, 2009). The read count was obtained with the phyton script *sam2counts* (https://github.com/vsbuffalo/sam2counts). Differentially expressed (DE) transcripts were identified using the edgeR package for R, the transcripts were normalized by cpm and filtered; 2 cpm in at least 2 samples (Robinson et al., 2010). Each sample was from a single seedling in a different experimental replicate. The samples were basically grouped by tissue and nutritional condition (Figure S2). For the tissue-treatment interaction only the transcripts with FDR < 0.05 and the three experimental replicates with the same expression sense than the final logFC (positive or negative) were considered as differentially expressed (DE). For the tissue analysis same parameters have been considered. The DE transcripts were used to construct a gene co-expression network. An unsigned network has been carried out using the R package WGCNA soft-thresholding power value of 9 (Langfelder & Horvath, 2008). From the network, the 10% of the transcripts with more connections in each module were considered hub genes.

These RNA-Seq data have been deposited in the NCBI’s Gene Expression Omnibus (Edgar et al., 2002) and are accessible through GEO Series with the accession number GSE175587 (https://www.ncbi.nlm.nih.gov/geo/query/acc.cgi?acc<u>= GSE175587</u>). Additionally, RNA-seq and network results are accessible through a database in html format that can be installable with R packages and downloaded from GitHub (https://github.com/ceslobfer/Rootapp).

Total RNA (1 ng) was retrotranscribed and amplified to verify the expression analyses for several DE genes by RT-qPCR. The cDNA synthesis and amplification protocol was carried out using the Conifer RNA Amplification (CRA+) protocol previously described by Cañas et al. (2014). Full step protocol is described in Methods S1. The amplification process was monitored using the ERCC RNA Spike-in kit (Thermo Scientific, Waltham, MA, USA) according to manufacturer’s instructions. The primers used for cDNA synthesis and amplification are listed in the Table S2.

### Functional annotation and enrichment analyses

The assembled transcriptome was functionally annotated with BLAST2GO (Götz et al., 2008) using DIAMOND software with *blastx* option (Buchfink et al., 2015) against the NCBI’s plants-*nr* database (NCBI Resource Coordinators, 2016). Blast results were considered valid with e-value < 1.0E-6. Singular enrichment analysis (SEA) of the GO terms was made in the AGRIGO v2.0 web tool under standard parameters using as GO term reference the whole assembled transcriptome annotation (Tian et al., 2017). Representative enriched GO was determined using REVIGO with 0.5 as dispensability cutoff value (Supek et al., 2011).

### RT-qPCR

The cDNA synthesis was performed using 1 μg of total RNA and iScript™ cDNA Synthesis Kit (Bio-Rad, Hercules, CA, USA) following manufacturer’s instructions. The qPCR primers were designed following the MIQE guidelines (Bustin et al., 2009). The primers are listed in the Table S2. qPCRs were carried out using 10 ng of cDNA and 0.4 mM of primers and 2X SsoFast^TM^ EvaGreen® Supermix (Bio-Rad, Hercules, CA, USA) in a total volume of 10 μL. Relative quantification of gene expression was performed using thermocycler CFX 384™ Real-Time System, Bio-Rad, Hercules, CA, USA). The qPCR program was as follows: 3 min at 95 °C (1 cycle), 1 s at 95 °C and 5 s at 60 °C (50 cycles) and a melting curve from 60 to 95 °C, to generate the dissociation curve to confirm the specific amplification of each individual reaction. The analyses were carried out as described by Cañas et al. (2014) using the MAK3 model in the R package *qpcR* (Ritz & Spiess, 2008). Normalization for gene expression of experimental design viability was performed using geometric mean of two reference genes, a *Saposin-like aspartyl protease* (pp_199988) and *Myosin heavy chain-related* (pp_58489) that were previously tested for maritime pine (Granados et al., 2016). Normalization for gene expression of LCM isolated tissues was performed using as reference gene a *SKP1/ASK1 family protein* (pp_18128) that was previously tested for maritime pine LCM samples (Granados et al., 2016). For the RT-qPCR analysis, three technical replicates of each sample and three biological replicates were made.

### Imaging of phytohormones in roots sections

The main phytohormones were localized in root apex cuts using nano-particle assisted laser desorption/ionization mass spectrometry imaging (Nano-PALDI-MSI). The elongation zones, which corresponded to the region from 2.5 - 5 mm of the root apex, were embedded in super cryoembedding medium (SCEM, Leica Biosystems, Wetzlar, Germany) and frozen in liquid nitrogen. The specimen block was cut into 10 µm sections using a cryostat (NX-70, Thermo Scientific, Waltham, MA, USA) set at −23°C in the chamber and at −25°C on the object holder. The sections were gently mounted on slides coated with indium tin oxide (ITO) (Bruker Daltonik GmbH, Billerica, MA, USA). Optical images of the sections were obtained by a virtual slide scanner (Nanozmmer-SQ, Hamamatsu Photonics, Shizuoka, Japan) before analysis by Nano-PALDI-MSI.

For Nano-PALDI-MSI, iron oxide-based nanoparticles (Fe-NPs) were prepared by stirring aqueous solutions of FeCl_2_·4H_2_O (5 mL, 100 mM; FUJIFILM Wako Pure Chemical), and 3-aminopropyltriethoxysilane (5 mL; γ-APTES; Shin-Etsu Chemical, Tokyo, Japan) at room temperature for 1 h. The resulting precipitate was washed several times with ultrapure water, resuspended in methanol (Moritake et al., 2007).

One milligram of Fe-NPs was resuspended in 1 mL methanol and sprayed on pine root tissue sections on ITO-coated glass slides with an airbrush (nozzle caliber, 0.2 mm). To obtain images, each data point on the section were irradiated with 200 laser shots in the positive ion detection mode of the mass spectrometer. Only signals between 80 and 800 *m/z* were analyzed to detect the corelated IAA (*m/z* 176.3), cytokinin (*m/z* 221.3), ACC (*m/z* 102.8), salicylic acid (*m/z* 139.4), JA (*m/z* 211.0) and ABA (*m/z* 265.0) as protonated ions, respectively. For each section, approximately 101,800 data points were obtained, 5 μm apart. The MS image was reconstructed from the obtained MS spectra with a mass bin width of m/z ± 0.1 from the exact mass using flexImaging 4.0 (Bruker Daltonik GmbH). The accurate mass of the ions was used for image generation, and mass accuracy and root-mean-square error (RMSE) were automatically calculated by the imaging software to avoid false-positive signals (Shiono and Taira, 2020). Comparisons of MS images were derived from the relative intensity for each signal normalized by the highest intensity spot on the slide. The peak intensity value of the spectra was normalized by dividing them with the total ion current (TIC) to achieve semi-quantitative analysis between control- and NH_4_ - treated roots.

### Phylogenetic analyses

Evolutionary analyses were performed in MEGA7 (Kumar et al., 2016). The protein sequence alignment was made with Muscle (Edgar, 2004). The evolutionary history was inferred using the Neighbor-Joining method (Saitou & Nei, 1987). The bootstrap consensus tree inferred from 1000 replicates is taken to represent the evolutionary history of the taxa analyzed (Felsenstein, 1985).

## RESULTS

### Tissue-specific transcriptomic response to NH_4_^+^ supply

In this study, tissue-specific transcriptome changes triggered by NH_4_^+^ nutrition were analyzed in the maritime pine root apex by a combination of laser capture microdissection and high-throughput RNA sequencing. Prior proceeding with tissue isolation by LCM, each experimental replicate was validated through the expression analysis of transcripts that were expected to be upregulated by NH_4_^+^ supply at 24 h post-irrigation (Figure 1) (Canales et al., 2011; Ortigosa et al., 2021). As expected, NH_4_^+^ induced the accumulation of transcripts coding for glutamine synthetase 1b (*PpGS1b*), antimicrobial peptide 1 (*PpAMP1*) and S-adenosyl methionine synthase (*PpSAMS*). Although not significant, the NADH glutamate synthase (*PpNADH-GOGAT*) expression also increased with NH_4_^+^ supply.

As described in the Materials and methods section, root apexes from seedlings that were irrigated with 3 mM NH_4_^+^ and harvested at 24 h were used for the isolation of four different tissues by LCM, namely, RC, RM, RDC and RDV (Video S1). The differential expression results are shown in Dataset S1. A total of 295 DE transcripts were identified in the low-input RNA-seq analysis (Figure 2A,B), of which 182 DE transcripts were downregulated (Figure 2A) and 113 were upregulated (Figure 2B). Among the isolated tissues, RDC showed the highest number of DE transcripts (107 upregulated and 70 downregulated) while RDV had the lowest response when seedlings were treated with NH_4_^+^ (12 upregulated transcripts). Interestingly, only 2 genes were upregulated in all root tissues, namely, two splicing isoforms from a common gene with unknown function.

**Figure 2.**
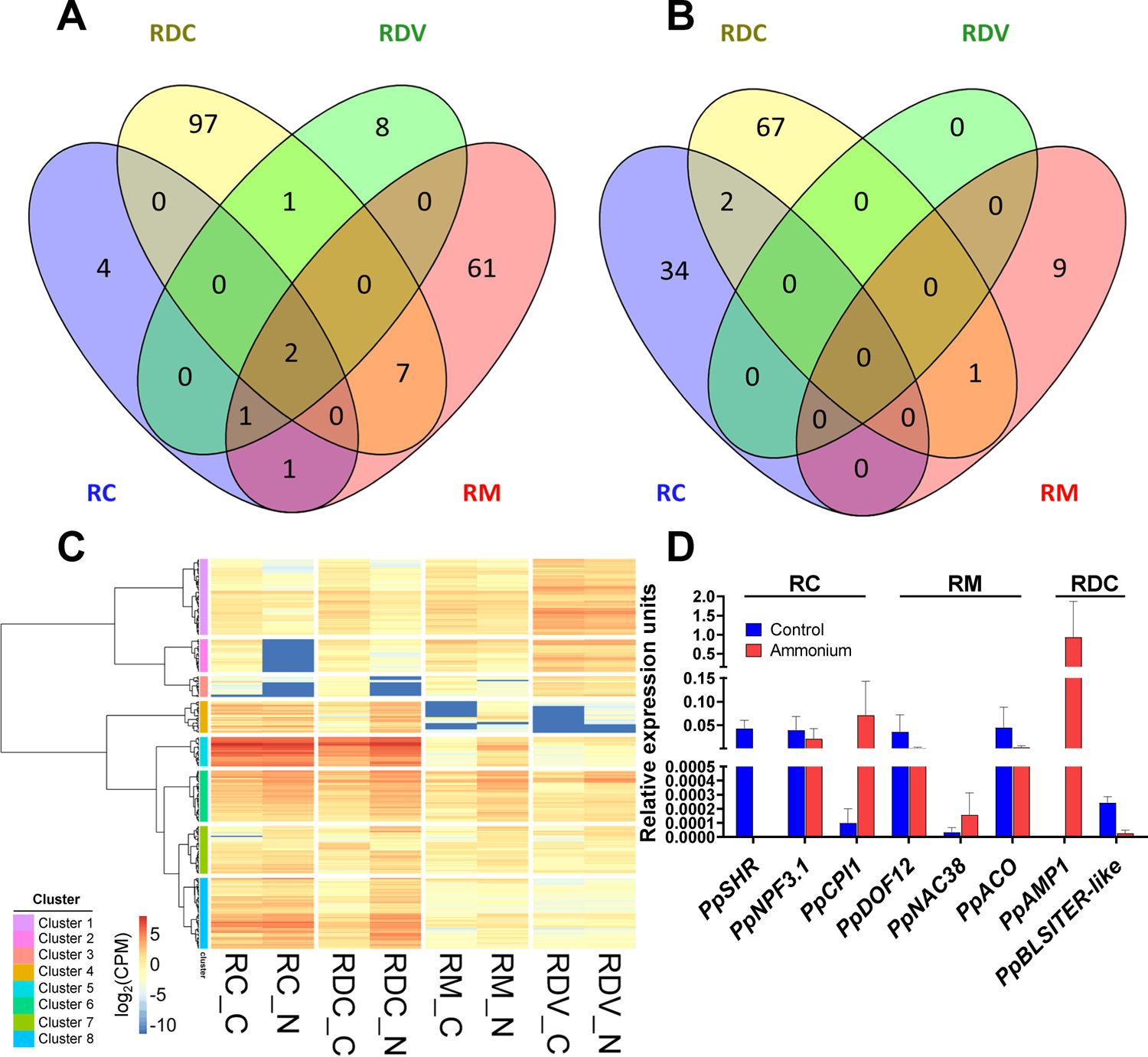
Low-input RNA-seq results in the different tissues of maritime pine root tip. (**A**) Venn diagram of down-regulated DE transcripts. (**B**) Venn diagram of up-regulated DE transcripts. (**C**) Heatmap and hierarchical clustering of DE transcripts. Expression values correspond to median CPM between biological/experimental replicates. The hierarchical clustering was made employing the Ward’s minimum variance method. (**D**) Experimental validation of low-input RNA-seq results for 8 DE transcripts through RT-qPCR. RC, the root cap; RM, root meristem; RDC, root developing cortex; and RDV, root developing vessels. *SHORT-ROOT* (*PpSHR*); *nitrate transporter 1 / peptide transporter family 3.1* (*PpNPF3.1*); *cycloeucalenol cycloisomerase* (*PpCPI1*); *DOF transcription factor 12* (*PpDOF12*); *NAC transcription factor 38* (*PpNAC38*); *aminocyclopropane-1-carboxylic acid oxidase* (*PpACO*); *antimicrobial peptide 1* (*PpAMP1*); *BLISTER-like transcription factor* (*PpBLISTER-like*).

The analysis of the DE transcripts revealed 8 expression patterns (Figure 2C). Among the groups with the most significant differences, the cluster 2 had transcripts repressed in RC by NH_4_^+^, and it contained several TFs, such as *PpSHR*, *PpDOF11* and *PpGATA20-like,* and transcripts related to phytohormones, such as *PpVAN3/FKD2* (Figure 2C; Dataset S2). In cluster 3, transcripts related to root development, such as *PpBAM1*, and phytohormones such as *flavanone 3-dioxygenase*, were found to be repressed by NH_4_^+^ in RC and RDC (Figure 2C; Dataset S2). In RDC and RM, NH_4_^+^ induced the transcripts framed into cluster 4, which are mainly related to defense, such as chitinases of class I and IV (Figure 2C; Dataset S2).

To validate our transcriptomic analysis, the expression value of several DE transcripts was corroborated by RT-qPCR (Figure 2D; Dataset S1). The downregulation of *PpSHR* (pp_346622) and *PpNPF3.1* (pp_58258) in RC; *PpDOF12* (pp_85381) and *PpACO* (pp_202153) in RM; and *PpBLISTER-like* (pp_170664) in RDC was confirmed. The upregulation of transcripts coding a *cycloeucalenol cycloisomerase* (*PpCPI1*; pp_189071) in RC and *PpAMP1* (pp_580007) in RDC was also corroborated.

### Functional enrichment analysis of tissue-specific transcriptomic response to NH_4_^+^ supply

The functional study of DE transcripts, including the SEA results, is summarized in Figure 3. In the most responsive tissues (RC, RM and RDC), NH_4_^+^ induced a general alteration in the amounts of transcripts coding TFs involved in development (most of them downregulated) and transcripts related to different phytohormones (Figure 3A-C). In the less responsive tissue (RDV) but also in RM and RDC, NH_4_^+^ promoted the accumulation of defense-related transcripts (Figure 3B-D). In the SEA results, no significant GO terms were found for DE transcripts upregulated in RC and RDV and downregulated in RM and RDV.

**Figure 3.**
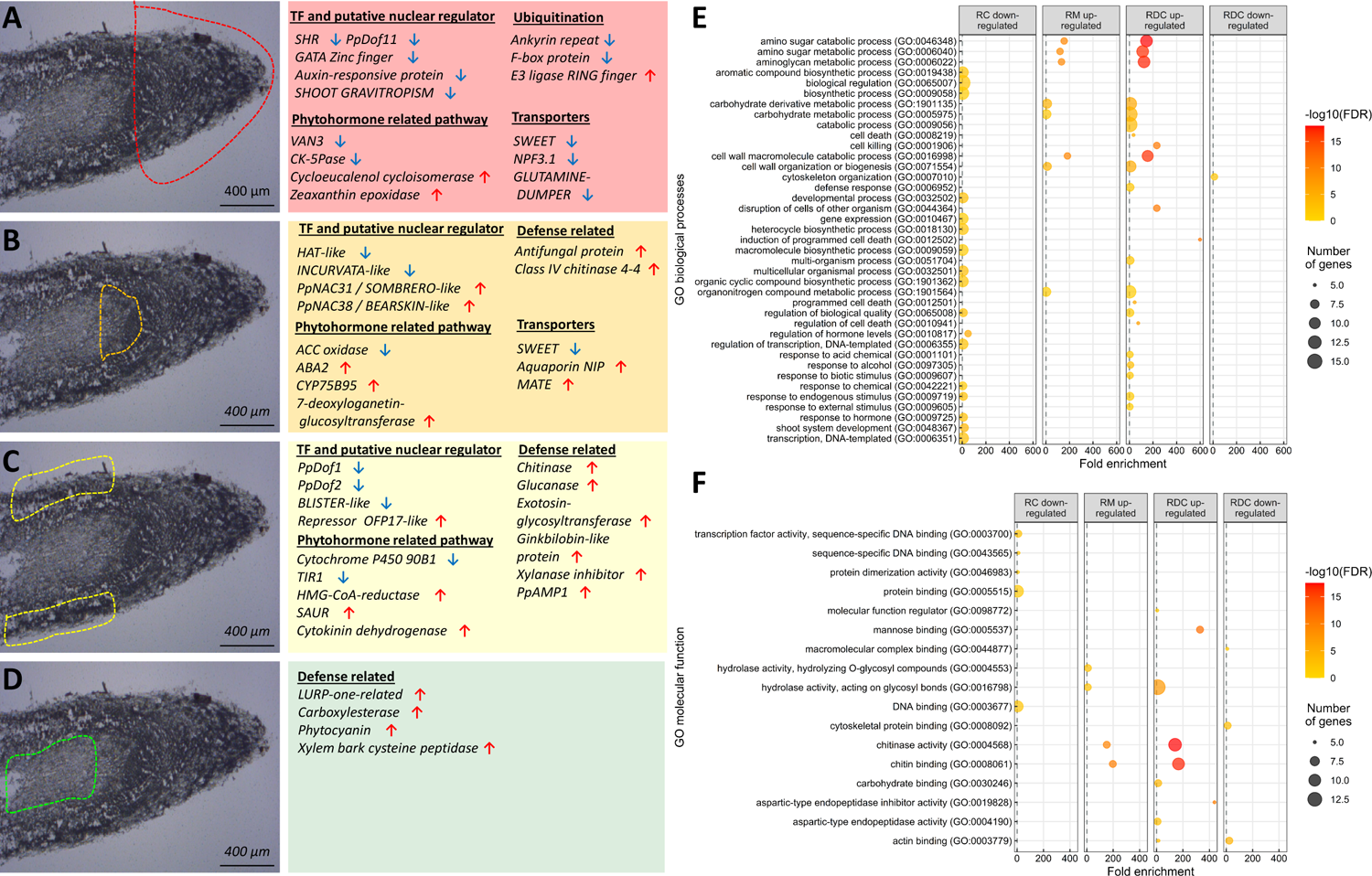
Resume of the main functional results obtained in the low-input RNA-Seq analysis from LCM isolated samples. Functions and DE transcripts in (**A**) the root cap (RC), (**B**) root meristem (RM), (**C**) root developing cortex (RDC) and (**D**) root developing vessels (RDV). Blue arrows indicate downregulated transcripts, red arrows upregulated ones. (**E**) Significant GO terms from Biological Processes category after a SEA analysis. (**F**) Significant GO terms from Molecular Function category after a SEA analysis.

The complete results of the SEA analyses are shown in Dataset S3. The largest number of SEA results were for transcripts downregulated in RC and upregulated in RDC. The downregulated transcripts in RC were significantly enriched in GO terms in the biological process (BP) category, such as “biological regulation” (GO:0065007), “gene expression” (GO:0010467), “response to hormone” (GO:0009725), “regulation of hormone levels” (GO:0010817) and “regulation of transcription, DNA-templated” (GO:0006355) (Figure 3E). The terms enriched in the molecular function (MF) category included “transcription factor activity” (GO:0003700), “protein binding” and “DNA binding” (GO:0005515 and GO:0003677) (Figure 3F). In the BP category for the downregulated transcripts in RDC only the “cytoskeleton organization” term (GO:0007010) was significantly enriched; however, for the upregulated transcripts in this tissue, numerous enriched functions were significant, such as those involved in defense responses such as “induction of programmed cell death” (GO:0012502), “cell killing” (GO:0001906), “defense response” (GO:0006952) and “cell wall macromolecule catabolic process” (GO:0016998) and “amino sugar catabolic process” (GO:0046348) (Figure 3E). Regarding the MF category for the upregulated transcripts in RDC, there were significant GO terms related to the response to biotic stress, such as “chitinase activity” (GO:0004568), “chitin binding” (GO:0008061), “mannose binding” (GO:0005537) and “hydrolase activity, acting on glycosyl bonds” (GO:0016798) (Figure 3F). Interestingly, the SEA results of the upregulated transcripts in RM were similar but limited for defense responses, which was also found for the upregulated transcripts in RDC (Figure 3E,F). Finally, the only enriched GO terms for the cellular component category were “extracellular region” (GO:0005576) and “extracellular space” (GO:0005615) for the transcripts upregulated in RDC tissue (Dataset S3).

### RNA-seq analysis in function of the root tissue

Although it has been studied more in deep in a previously work (Cañas et al., 2017), the transcriptional changes underlying different tissues that form the root apex of maritime pine (RC, RM, RDC and RDV) have been briefly analyzed (Dataset S1). The comparison of the DE transcripts showed a set of characteristic genes of each tissue. This gene core was composed by 1014, 2, 529 and 2692 DE transcripts for RC, RM, RDC and RDV, respectively (Figure S3A). The functional study of the core DE transcripts revealed the main functions for these tissues (Dataset S3). Some of the enriched GO terms for RC are “aminoglycan metabolic process” (GO:0006022), “cell wall organization or biogenesis” (GO:0071554), “dehiscence” (GO:0009900) or “polysaccharide catabolic process” (GO:0000272) (Figure S3B). For RDC gene core, GO terms such as “anatomical structure formation involved in morphogenesis” (GO:0048646), “catabolic process” (GO:0009056) or “response to stimulus” (GO:0050896) were observed (Figure S3B). On the other hand, RDV was the tissue that showed the highest number of DE transcripts when compared to the rest of the tissues, which presented enrichment GO terms such as “auxin influx” (GO:0060919), “carbohydrate metabolic process” (GO:0005975), “fluid transport” (GO:0042044) or “phloem or xylem histogenesis” (GO:0010087), among others (Figure S3B). Interestingly, in RM only 2 DE transcripts were identified as characteristic and both transcripts are splicing variants of the same gene encoding a bHLH TF, similar to the *Arabidopsis* FAMA TF (AT3G24140) (Figure S3C).

### Transcription factors affected by NH_4_^+^ in the apex of maritime pine roots

The DE transcripts were individually analyzed looking to identify different kinds of regulators, such as TFs and transcripts involved in the phytohormone response. From a total of 295 DE transcripts, 31 TFs were identified (Figure 4, Table 1, Dataset S1). All 10 FTs identified in RC were repressed in the presence of NH_4_^+^. However, in RM and RDC, 6 and 4 TF transcripts were downregulated while 5 and 6 TF transcripts were upregulated, respectively. Interestingly, a high abundance of TFs related to root growth and development was observed (Table 1), such as the strong downregulation of *PpSHR* in RC, *INCURVATA-like* and homeobox-leucine zipper protein *HAT-like* both in RM and involved in meristem developmental regulation, and the TF identified as *zinc finger C2H2 SHOOT GRAVITROPISM* in RC related to the gravitropism response. *PpSHR* was the most highly repressed TF (−11 logFC). Furthermore, the induction of two transcriptional repressors of different plant developmental processes was observed, with *PpNAC31* identified as an *SMB-like* NAC TF in RM and an *OFP17*-like repressor in RDC. In addition, novel TFs in maritime pine were identified. Two of them belong to the DOF-family and were named as *PpDOF11* and *PpDOF12*, and they were both downregulated in RC and RM, respectively. Phylogenetic analysis of these two new DOF-type TFs revealed that *PpDOF11* is grouped with *AtDOF1.4* (AT1G28310) and *OsDOF7.2* (LOC_Os07g32510) into subfamily E and that *PpDOF12* is grouped with members of subfamily A of *P. pinaster*: *PpDOF4*, *PpDOF7*, *PpDOF8* and *PpDOF10* (Figure S4, Table S3). Additionally, a member of the NAC family (*PpNAC38)* that was upregulated in RM tissue was identified. This TF has homology to *BRN1* (AT1G33280) (e-value: 7e-96) and *BRN2* (AT4G10350) (e-value: 3e-97) of *Arabidopsis thaliana* and is framed within the NAC subfamily C (Figure S5, Table S4).

**Figure 4.**
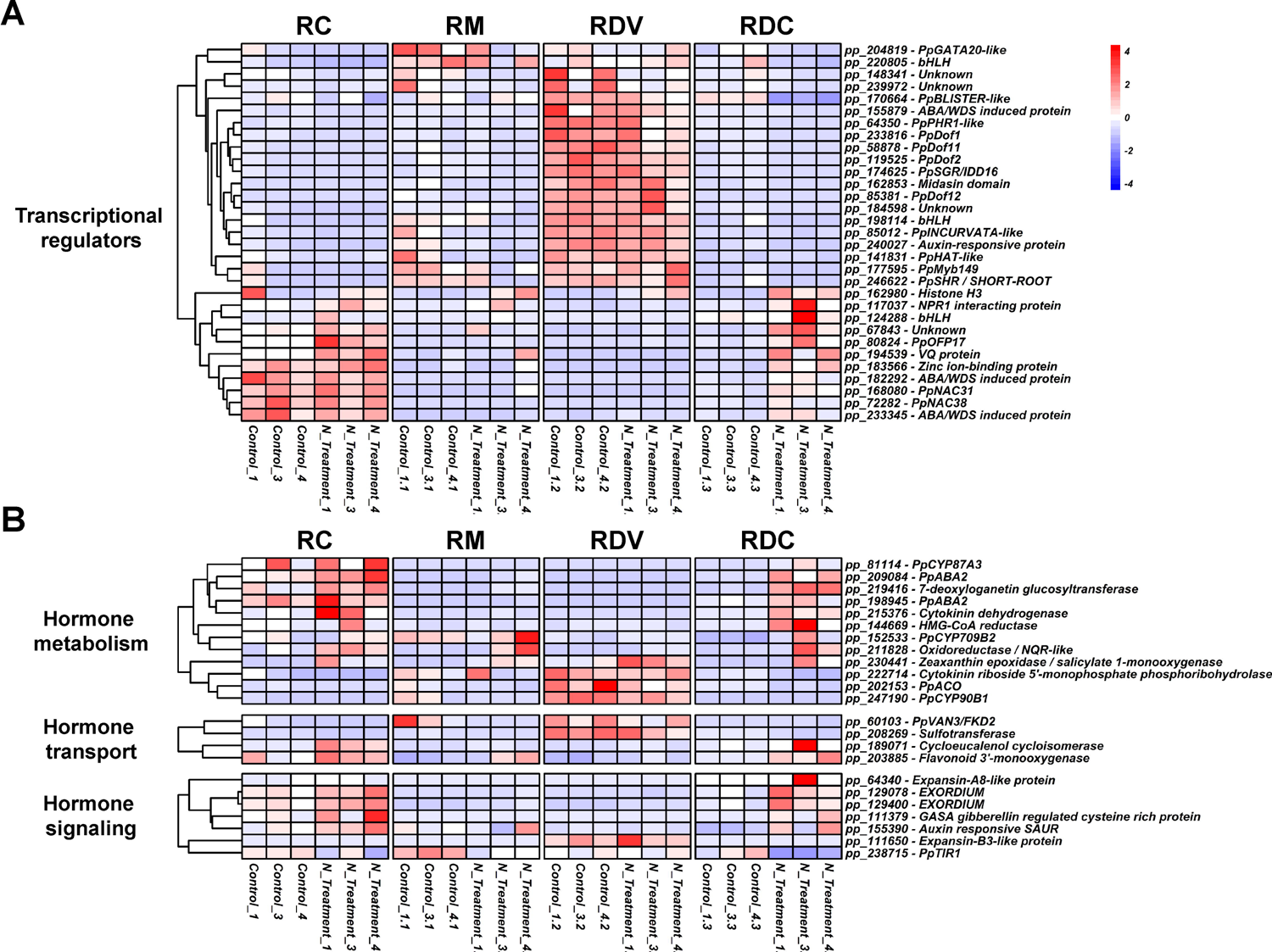
Expression heatmaps of differentially expressed transcriptional regulators and hormone related transcripts. (A) Transcription factors and other putative regulators. (B) Transcripts involved in hormone metabolism, transport and signaling. CPM expression values were normalized by transcript.

A gene co-expression network was constructed to identified hub TFs and putatively determine the processes controlled by them. Thirteen transcripts were identified as hubs corresponding to 8 different TFs: *PpDOF1*, *PpDOF2*, *PpDOF11*, *PpGATA20-like*, *PpINCURVATA-like*, *PpMyb149*, *PpSGR/IDD16* and *PpSHR*. All of them were DE for NH_4_^+^ treatment in different tissues except RDV but all of them were more expressed in RDV than in the rest of tissues except *PpGATA20-like* (Dataset S1). The functions and expression localization of the most correlated transcripts (>|0,9|) were analyzed (Figure S6; Dataset S4). Most of the transcripts were correlated with *DOF* TFs, *PpINCURVATA-like* and *PpSGR/IDD16* but also belonged to the RDV gene core (1879 from 2362) (Figure S6A). Thus, the enriched functions were mainly related with RDV functions such as “auxin influx” (GO:0060919), “carbohydrate metabolic process” (GO:0005975), “fluid transport” (GO:0042044) or “phloem or xylem histogenesis” (GO:0010087) (Figure S6B; Dataset S5). Interestingly, transcripts correlated with *PpGATA20-like* were enriched in functions such as “gene expression” (GO:0010467) and “ribosome biogenesis” (GO:0042254) (Figure S6B; Dataset S5).

### Phytohormone-related genes

Twenty-six DE transcripts related to phytohormone pathways were identified (Table 1), of which 19 were upregulated and 7 were downregulated (Figure 4, Dataset S1). Most of these DE transcripts were related to IAA (Table 1). Some interesting examples are the downregulation of transcripts encoding proteins required for PIN transporters localization, such as VAN3/FKD2 Auxin canalis/PH2 (pp_60103) in RC and the upregulation of a *PpCPI1* (pp_189071) in RC, which is related to PIN transporters endocytosis. In addition, in RDC, the repression of transcripts coding for the IAA receptor TIR1/AFB (pp_238715) and for a sulfotransferase (pp_208269) with a high similarity degree to SULT202B1 (AT3G45070) (e-value: 5e-72) was observed.

CK-related genes were the second most represented phytohormone-related transcripts, and strong repression of CK riboside 5’-monophosphate phosphoribohydrolase (pp_222714) (−6.9 logFC), a CK-activating enzyme, was observed in RC. However, upregulation of a transcript coding for a CK dehydrogenase (pp_215376), a CK-inactivating enzyme, was also observed in RDC. The repression of transcripts related to brassinosteroids and ethylene (ET) biosynthesis, e.g., cytochrome P450 90B1 (pp_247190) in RDC and *PpACO* (pp_202153) in RM was also observed. In addition, transcripts related to the gibberelin (GA) response and abscisic acid (ABA) biosynthesis were upregulated, such as a GASA gibberellin-regulated cysteine-rich protein (pp_111379) in RDC and two ABA2 coding transcripts that might be involved in ABA biosynthesis (pp_198945, and pp_209084) in RM and RDC.

### Phytohormone detection in maritime pine root apex

To corroborate whether NH_4_^+^ nutrition altered the spatial allocation of several phytohormones as the transcriptomic data suggest, the distribution of multiple phytohormones was determined by Nano-PALDI-MSI in the apex of maritime pine roots (Shiono & Taira, 2020). In comparison to control seedlings, changes in the patterns of IAAs, CK (tZ, *trans*-Zeatin), ACC, ABA, salicylic acid (SA) and jasmonic acid (JA) were observed in the presence of NH_4_^+^ (Figures 5 and S5). In the control plants, an IAA gradient was observed, the maximum level was detected in the most distal area of the principal root apex (Figure 5A); however, in the NH_4_^+^-treated roots, the IAA maximum was detected in a more distant zone from the root tip (Figure 5A). CK (tZ) showed a wide distribution in the roots of control plants (Figure 5B). In the NH_4_^+^-treated seedlings, CK was distributed in the outer tissues of the root mainly below the root tip (Figure 5B). Regarding ET, its precursor ACC was found in a very restricted area of the root tip in control and NH_4_^+^-treated plants, showing no changes when plants were supplied with NH_4_ (Figure 5C). ABA was detected mainly in the RDC tissue in both control and treated seedlings, and it was higher under NH_4_^+^ supply and distributed below the root apex (Figure 5D). SA and JA phytohormone distributions were also affected by the presence of NH_4_^+^ (Figure S7). SA was located at the end of the root tip, while JA was also detected in the elongation zone in control plants. When the seedlings were supplied with NH_4_, SA showed a vaguer spatial distribution with respect to the observed distribution in the control plants (Figure S7A) while JA tended to slightly increase in the outermost parts of the root apex ends and disappeared from the elongation zone (Figure S7B).

**Figure 5.**
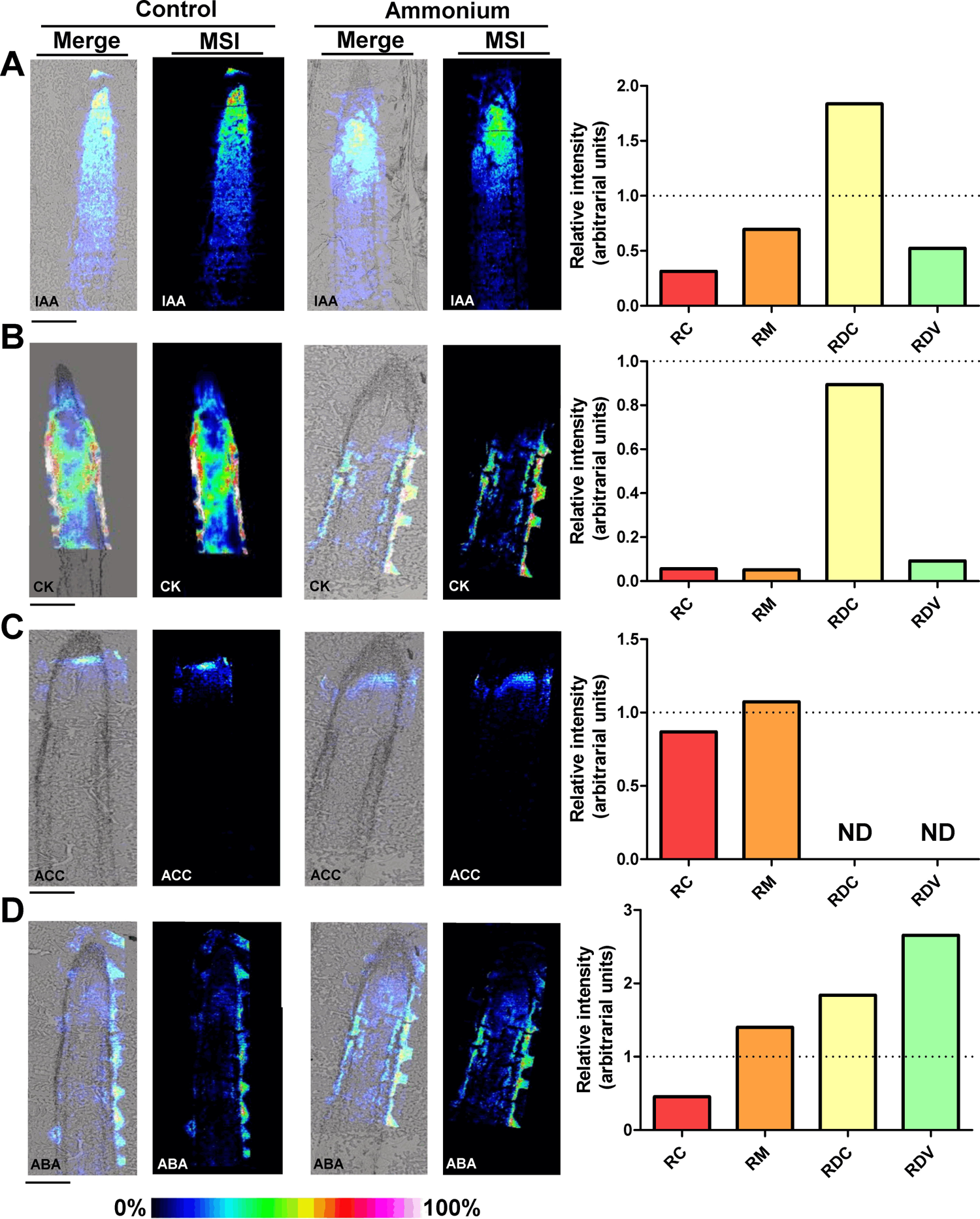
Phytohormone localization in root apex sections. The main phytohormones were localized in root apex cuts of control and 3 mM NH_4_^+^ treated seedlings. The phytohormone identification and imaging was made using nano-particle assisted laser desorption/ionization mass spectrometry imaging (Nano-PALDI-MSI). (**A**) Auxin / Indole-3-acetic acid (IAA); (**B**) cytokinin (CK); (**C**) aminocyclopropane-1-carboxylic acid (ACC); (**D**) abscisic acid (ABA). Size bars correspond to 500 µm. Column graphs show relative hormone accumulations respect to control samples in the approximate LCM areas. Dashed lines mark hormone accumulation in the control roots.

### Maritime pine root phenotype

Maritime pine root morphological studies showed no differences between treatments regarding whole root biomass accumulation (Figure 6A). When the weight of the principal roots (PRs) was measured, NH_4_^+^-fed seedlings exhibited a significant reduction in PR weight compared to water- and NO_3_^-^-treated plants (Figure 6B) and a statistically higher root length when pine seedlings were supplied with either inorganic nitrogen form (Figure 6C). Moreover, NH_4_^+^ promoted an increase in LR number and weight compared to the water and NO_3_^-^ treatments (Figure 6D,E), although no statistically significant differences were observed regarding LR density (LRD) between treatments (Figure 6F).

**Figure 6.**
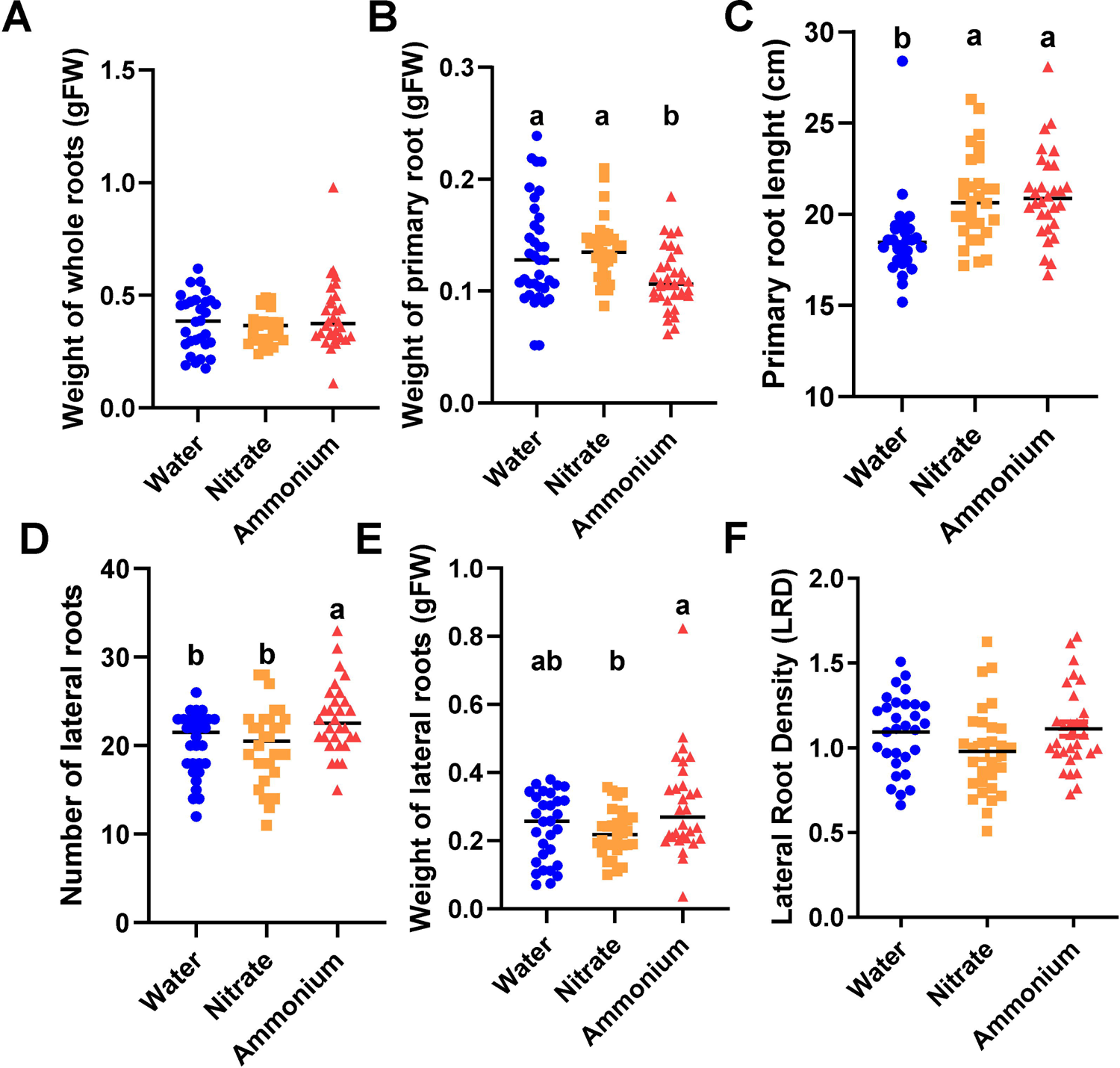
Root growth and architecture parameters. Pine seedlings were cultivated for 74 days and irrigated once a week with 50 mL of water, 3 mM KNO_3_ and 3 mM NH_4_Cl, respectively. Six different parameters were measured after harvesting: (**A**) weight of whole roots; (**B**) weight of primary root; (**C**) primary root length; (**D**) number of lateral roots; (**E**) weight of lateral roots; (**F**) lateral root density. Significant differences were determined with a one-way ANOVA. Letters above the conditions show significant differences based on a Turkey post-hoc test (*P* < 0.05) with n = 30.

## DISCUSSION

Despite the obvious environmental and economic importance of conifers, little is known about their response to diverse nutrients at the molecular and developmental levels. Since they represent an ancient lineage of gymnosperms, conifers are of particular interest from an evolutionary perspective because they have common but also differential responses with angiosperm models. The aim of the present work was to explore the early response of maritime pine to NH_4_^+^ nutrition in the different tissues of the root apex. For this purpose, tissue isolation was performed by LCM and combined with low-input RNA-seq to determine the local transcriptomic response in the growth area of the root. This strategy avoids the dilution effect of transcripts with low and much localized expression, revealing new genes not previously annotated (Cañas et al., 2017), such as *PpDOF11-12* and *PpNAC38*. Clearly a dilution effect can be observed when compared the expression of marker genes in the whole roots (Figure 1) and in the specific tissue transcriptomics (Figure S8A). Supporting this, from 47,118 transcripts used in the tissue specific RNA-seq 8,491 transcripts were not previously identified in whole root transcriptomic experiments (Dataset S1). A similar number of DE transcripts was obtained in the present work (295) when compared to the same NH_4_^+^ nutrition treatment over the whole root (350) (Ortigosa et al., 2021) (Figure S8B). However, only 3 transcripts with the same identifiers are shared, although there are different transcripts from the same gene that are DE in both analyses, for example *PpAMP1* (RDC - pp_58007 / whole root - pp_58005) (Figure S8C).

The response to NH_4_^+^ in RDC involved the expression of defense-related genes as previously described in maritime pine and other model plants (Figure 3C,E,F) (Canales et al., 2010; Patterson et al., 2010; Ravazzolo et al., 2020; Ortigosa et al., 2021). Obviously, this response might be linked to the peripheral localization of RDC in the roots since the defense proteins encoded by the expressed transcripts develop their functions in the extracellular media (Dataset S3). Some examples are the upregulation of different *ginkbilobin-like*/*embryo abundant protein* transcripts (e.g. pp_252647), *chitinases* (e.g. pp_117806) and the *PpAMP1* (pp58007), which are consistent with results previously reported in maritime pine roots (Canales et al., 2010; Ortigosa et al., 2021).

In angiosperm model plants, NH_4_^+^ alters RSA by inhibiting root elongation, stimulating LR branching and affecting root hair development (Y. Liu & von Wirén, 2017). RSA principally targets root system processes, such as root elongation, root gravitropism and LR branching, appearing to occur in the root tip (Li et al., 2014). Based on the presented transcriptomic analyses, early root exposure to NH_4_^+^ caused a wide impact on the expression of TFs related to root growth and development in pine (Figure 2, Table 1). Thus, NH_4_^+^ caused a strong repression of *PpSHR* (pp_246622) in RC tissue, a GRAS-type TF involved in the regulation and coordination of root development, including phloem differentiation (Kim et al., 2020). Interestingly, in RDC tissue, it was observed that *PpBAM-like* (pp_19744) transcripts were also severely affected by NH_4_^+^ (Dataset S1), which is consistent with previous cell-specific transcriptomic profiles observed in *Arabidopsis* (Brady et al., 2007). BAM1/2 kinase receptors are required for SHR-dependent formative divisions in roots and for the proper CYCD6;1 expression, which is required to promote division in cortex endodermal initial daughter cells (Crook et al., 2020).

Additionally, a progressive reduction in the abundance of IAA efflux carriers (PINs) in *Arabidopsis shr* mutants has been described (Lucas et al., 2011), and NH_4_^+^ negatively affects the expression of PIN2 and AUX1 coding transcripts (Y. Liu et al., 2013). The expression of genes coding for PIN transporters was not affected in pine roots. Recently, it has been linked IAA signaling pathway to acidification (Jia et al., 2020; Meier et al., 2020).

However, based on our results, no significant changes were observed in the expression of pH-related genes. However, it was observed that the expression of coding transcripts for proteins involved in IAA transporter localization and components of the IAA signaling pathway were affected by NH_4_^+^. This is the case for *PpCPI1* (pp_189071), which is upregulated in RC and required for PIN2 endocytosis (Men et al., 2008), and the strong repression of *VAN3/PH2/FKD2* (pp_60103) in RC and RM tissues (Table 2), which is crucial in *Arabidopsis* for the vascular leaf pattern formation by the proper PIN transporter localization (Hou et al., 2010). These results suggest that NH_4_^+^ triggers an alteration in polar auxin transport (PAT) involving a highly coordinated transcriptomic response between tissues of the pine root. The Nano-PALDI-MSI data confirmed that under a supply of NH_4_^+^, PAT was impaired, thus promoting a putative local increase in IAA presumably in RC and/or RM tissues where cambial development takes place based on IAA-related transcripts and TF expression (Figures 5A and 7), which could suggest a different regulatory mechanism of those pH-dependent described in rice and *Arabidopsis* (Jia et al., 2020; Meier et al., 2020). One possible alternative for this PAT alteration could be IAA conjugation with sugars/amino acids as previously described (Tamura et al., 2010; Di et al., 2021). However, our data revealed non-significant expression changes of IAA conjugating genes (Dataset S1). This suggests that the NH_4_^+^ concentration used in this work is not excessive for maritime pine compared with those applied before (>5 mM NH_4_^+^) in rice and *Arabidopsis* (Tamura et al., 2010; Di et al., 2021), which is supported by the non-inhibition of PR growth under NH_4_^+^ supply (Figure 6C). Therefore, other mechanisms could be acting during the observed PAT process in the root apex of maritime pine.

**Figure 7.**
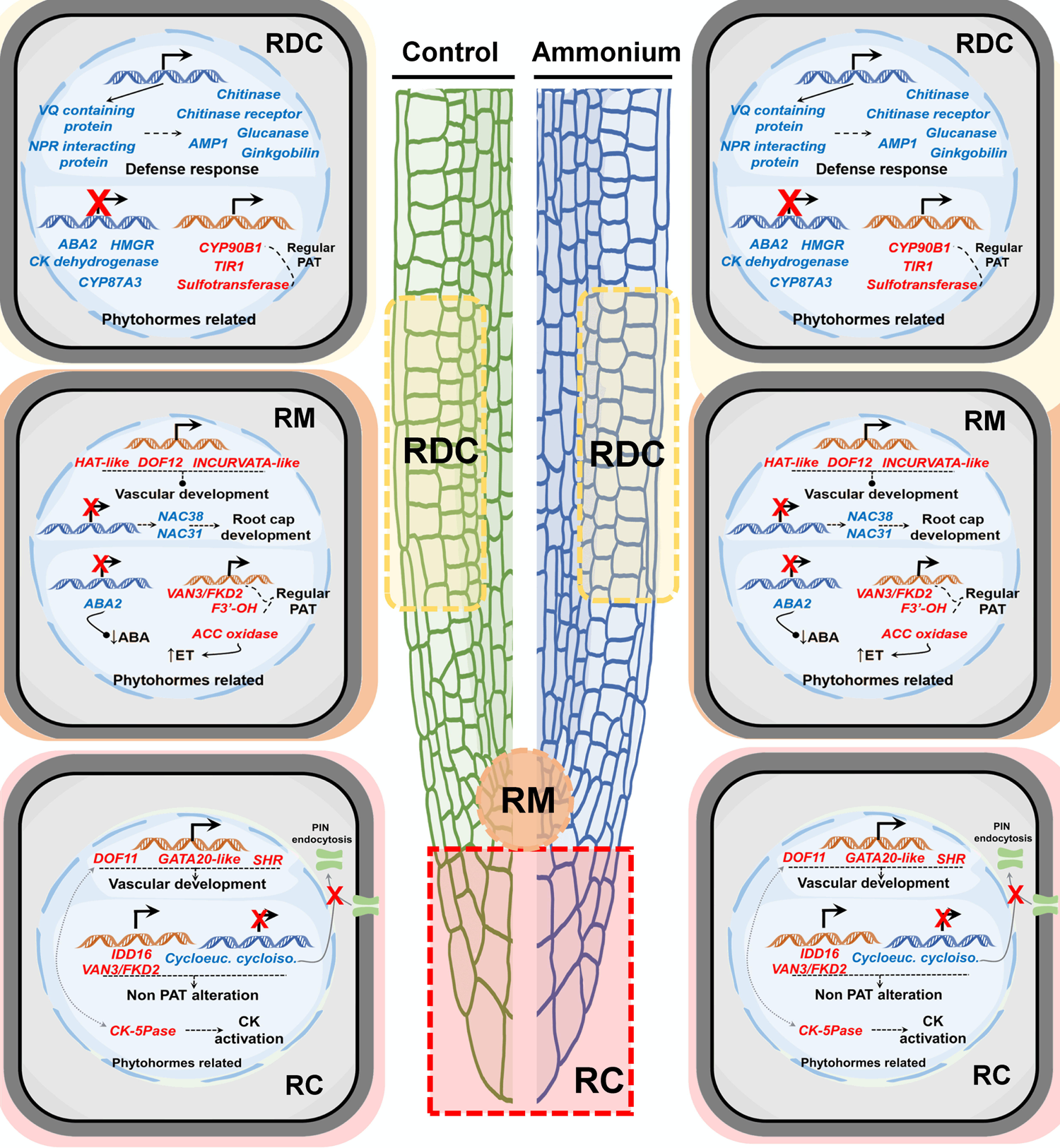
Schematic representation of the transcriptional response induced by NH_4_^+^ supply in maritime pine seedling roots 24 h after irrigation. Transcripts in red letters are upregulated. Transcripts in blue letters are downregulated. The root tip tissues are the root cap (RC) root meristem (RM) and root developing cortex (RDC). Results for root developing vessels (RDV) are not shown since they were very limited.

In addition, our results differ from those previously described in *Arabidopsis* where the presence of NH_4_^+^ does not seem to affect the distribution of IAAs in the root tip (Y. Liu et al., 2013). This finding could be related to root vascular development and root cell differentiation since adjustments to IAA flux are involved in root cambium development (Marhava et al., 2018), and SCR, a partner of SHR, is involved in the expression and polar localization of PINs (Xu et al., 2006).

Following this line of argument, it is interesting to mention that four maritime pine DOFs were significantly downregulated: *PpDOF11* (pp_58878) and *PpDOF12* (pp_85381) in RC and RM tissues respectively, and *PpDOF1* (pp_233816) and *PpDOF2* (pp_119525) in RDC (Figure 2), which were preferentially expressed in tissues close to vascular vessels (Figures S9-S12). Therefore, it is tempting to suggest that these DOFs may be involved in the development of root vessels, as described for PHLOEM EARLY DOF (PEAR) proteins, which are regulated by CK levels (Miyashima et al., 2019). This is also supported by the network analysis and the functions of the most correlated transcripts to these DOFs (Figure S6B). Interestingly, the CK distribution under NH_4_^+^ supply (Figure 5B) correlates with the repression of *CK riboside 5’-monophosphate phosphoribohydrolase* (pp_222714), which is a CK activator, in RC (Kuroha et al., 2009) and the upregulation of a *CK dehydrogenase* (pp_215376), which is involved in CK degradation (Werner et al., 2003), in RDC (Figure 3A,C). All these results suggest that NH_4_^+^ could promote an alteration of CK spatial distribution in maritime pine roots, which could be the reason for the wide repression of DOF TFs in RC and RM tissues and its possible role as regulators of cambium development (Figure 7). Phytohormone imaging showed that CK and IAA were distributed in different zones under a supply of NH_4_^+^ (Figure 5A,B), which was possibly due to the opposite roles of these phytohormones in the root meristem since CK negatively regulates QC specification and functions by modulating the IAA response (Zhang et al., 2013).

Additionally, homeobox-leucine zipper (HD-Zip) and NAC-domain TFs are described to take part during development processes (Ochando et al., 2006; Bennett et al., 2010). In *Arabidopsis*, several HATs play important roles in development including the regulation of meristem and procambial maintenance and/or formation (Prigge et al., 2005; Roodbarkelari & Groot, 2017). Accordingly, two HD-Zip coding transcripts identified as *HAT-like* (pp_141831) and *INCURVATA-like* (pp_85012) were downregulated in RM tissue (Table 1). These findings are consistent with the repression of TFs related to the root vascular development process and suggest that NH_4_^+^ could affect PR vascular development during early exposure (Figure 7). Nevertheless, long-term NH_4_^+^ nutrition promotes PR growth but causes a decrease in PR weight (Figure 6B,C), suggesting that root radial growth could be affected by this transcriptional network. In future studies, cell division and elongation should be analyzed in the root these zones. Unfortunately, the exact anatomy of root meristem with the quiescent center position has not yet been defined in conifers.

Interestingly, in RM, the upregulation of two NAC TFs, *PpNAC31* and *PpNAC38,* similar to the *Arabidopsis SMB* and *BRN* genes (Table 1, Figure S4), is consistent with the expression patterns of *SMB*, *BRN1* and *BRN2*, which largely overlap in *Arabidopsis* roots (Bennett et al., 2010). SMB represses stem cell-like divisions in the root cap daughter cells (Bennet et al., 2014; Kamiya et al., 2016), and together with BRN1 and BRN2, it regulates cellular maturation of the root cap in *Arabidopsis* (Bennett et al., 2010). These findings are consistent with the reduction in the number of root cap cells under NH_4_^+^, as described by Y. Liu et al., (2013), thus providing a possible explanation for this observation in *Arabidopsis*. In rice, the endogenous ABA content alleviates NH_4_^+^ rice toxicity, promoting NH_4_^+^ assimilation by stimulating the GS/GOGAT cycle and enhancing antioxidant activities (superoxide dismutase, ascorbate peroxidase and catalase) (Sun et al., 2020). In the maritime pine root apex under NH_4_^+^ supply, ABA levels increased (Figure 5D), and transcripts encoding ABA2 enzymes (e.g. pp_198956) involved in ABA biosynthesis were upregulated in RM and RDC, and several putative ABA-induced TFs (e.g. pp_233345) were upregulated in RC and RM (Dataset S1). However, the GS/GOGAT cycle was apparently not induced, although it was observed in the whole root response to NH_4_^+^ supply (Ortigosa et al., 2021) (Figure 1). In general, there was no expression changes in genes involved in N uptake and assimilation at the root apex of pine in response to NH_4_^+^ supply. In angiosperms, the upregulation of GS/GOGAT cycle has been described in specific tissues such as root epidermis around root hair zone in rice (Ishiyama et al., 1998; Ishiyama et al., 2004a) and *Arabidopsis* (Ishiyama et al., 2004b; Konishi et al. 2017). This raises the question of whether conifers might have a different response to NH_4_^+^ availability at the root apex than angiosperms.

Regarding antioxidant metabolism, different transcripts involved in ascorbate production were downregulated (Dataset S1), including *PpBLISTER-like* TF (pp_170664) in RDC tissue (Purdy et al., 2011) and two *L-gulonolactone oxidases* (pp_183087 and pp_234650) in RC and RDC tissues (Aboobucker et al., 2017). This finding suggests that the glutathione-ascorbate cycle is not operative in maritime pine roots exposed to NH_4_^+^, which is consistent with the levels of oxidized glutathione observed under long-term experiments (Ortigosa et al., 2020).

Finally, ET is involved in NH_4_-responsive pathways in rice (Sun et al., 2017) and in cambial development in poplar trees (Love et al., 2009). In *Arabidopsis,* the ET levels increased under a supply of NH_4_^+^, which reduced lateral root formation because of the repression of auxin transporter 1 (AUX1) (Li et al., 2014). In contrast, the results in maritime pine suggest a reduction in ET biosynthesis following NH_4_^+^ application. A transcript encoding an ACO was repressed in RM, which is consistent with proteomic and epitranscriptomic results in maritime pine whole roots (Ortigosa et al., 2021). This finding is consistent with the downregulation of *BLISTER-like* TF and *L-gulonolactone oxidases* since ascorbate is one of the ACO enzyme substrates (Dilley et al., 2013). However, ACC was not affected by NH_4_^+^ treatment (Figure 5C). Accordingly, NH_4_^+^ stimulated the LR components (number and weight) and PR length (Figure 5) in maritime pine, which is consistent with the increased number of root apexes under NH_4_^+^ nutrition compared to NO_3_^-^ nutrition in *Pinus massoniana* Lamb. (Ren et al., 2020). Taken together, the results suggest that maritime pine reduces ET biosynthesis under NH_4_ ^+^ supply, which could be linked to the PAT and CK alteration observed (Figure 4A,B) as well as to the expression patterns observed for TFs related to cambium development (Table 1) due to the importance of IAA-ET-CK crosstalk during root development (J. Liu et al., 2017).

## CONCLUSIONS

NH_4_^+^ nutrition is a complex process in plants since NH_4_^+^ is a nutrient that can be toxic when supplied in excess. However, conifers in general are quite tolerant to NH_4_^+^.

Therefore, the identifying of the differential mechanisms underlying NH_4_ tolerance in conifers would be enormously helpful for improving NH_4_ tolerance not only in these trees but also in other crops. In contrast to other plant models, supplying NH_4_^+^ to pine represses ET and ascorbate production and alters IAA and CK patterns, thus regulating a transcriptional network in the short term. Consequently, the growth and development of PR and LRs are stimulated in pine.

The expression profiles of genes related to root cambium development and phytohormones suggest a molecular mechanism underlying changes in the RSA phenotype that includes IAA-CK-ET crosstalk and a transcriptional network at least during the early root response to NH_4_^+^ supply. The results reported in the present study tentatively link *SHR*, and other TFs to NH_4_^+^ nutrition and its phenotypic effect on RSA, which likely affect early vascular development (Figure 7).

This work provides new and valuable data to unravel the mechanisms involved in the response of maritime pine to NH_4_^+^ nutrition. However, further research efforts are required to reach a full understanding of the molecular basis of NH_4_^+^ tolerance.

## ACKNOWLEDGEMENTS

Acknowledgments: This research was funded by Spanish *Ministerio de Ciencia e Innovación*, grant numbers BIO2015-73512-JIN MINECO/AEI/FEDER, UE, RTI2018-094041-B-I00 and EQC2018-004346-P. FO was supported by grants from the *Universidad de Málaga* (*Programa Operativo de Empleo Juvenil vía SNJG, UMAJI11, FEDER, FSE, Junta de Andalucía*) and BIO-114*, Junta de Andalucía*.

## Author Contributions

FO, HS and ST have performed the experiments; CLF has performed transcriptomic data analysis; FO and RAC have wrote the manuscript; CA and FMC made additional contributions and edited the manuscript; CA, FMC and RAC have acquired the funds; FO and RAC have planned and designed the research.

## Data Accessibility Statement

The data that support the findings of this study are available from different databases, supporting information and from the corresponding author upon reasonable request. RNA-Seq data have been deposited in the NCBI’s Gene Expression Omnibus and are accessible through GEO Series with the accession number GSE175587 (https://www.ncbi.nlm.nih.gov/geo/query/acc.cgi?acc= GSE175587). The transcriptome shotgun assembly project has been deposited at DDBJ/EMBL/GenBank under the accession GJFX00000000. The version described in this paper is the first version, GJFX01000000.

## Conflict of Interest

The authors have no conflict of interest.

## Supporting information

**Video S1.** LCM cutting procedure.

Table S1. LCM low-input RNA sequencing results.

Table S2. List of used primers.

Table S3. Protein sequences of DOF transcription factors in maritime pine, *Arabidopsis* and rice.

Table S4. Protein sequences of NAC transcription factors in maritime pine and *Arabidopsis*.

**Method S1.** Complete LCM and cDNA amplification procedures.

Figure S1. Aspect of root architecture after long-term nutritional treatments.

Figure S2. MDS plot of the LCM low-input RNA-seq samples.

Figure S3. RNA-seq differential expression analysis between tissues. (**A**) Venn diagrams shows the amounts of tissue-specific DE transcripts. (**B**) Functions of the transcripts that were always significantly expressed in a specific tissue. Significant GO terms from Biological Processes category after a SEA analysis with a 0.4 cut-off value for dispensability. (**C**) Expression of transcription factor PpFAMA-like (pp_179856) in the different isolated tissues of the root apex. PpFAMA-like was differentially expressed in the RM respect to the rest of tissues but not in the nutritional treatment comparison.

Figure S4. Phylogenetic tree of DOF transcription factor family in maritime pine, *Arabidopsis* and rice. The protein sequences employed to infer the phylogenetic tree are presented in the Table S3. The protein sequence alignment was made with Muscle (Edgar, 2004). The evolutionary history was inferred using the Neighbor-Joining method (Saitou and Nei, 1987). The bootstrap consensus tree inferred from 1000 replicates is taken to represent the evolutionary history of the taxa analyzed (Felsenstein, 1985). Branches corresponding to partitions reproduced in less than 50% bootstrap replicates are collapsed. The evolutionary distances were computed using the Dayhoff matrix-based method (Schwarz and Dayhoff, 1979) and are in the units of the number of amino acid substitutions per site. The analysis involved 78 amino acid sequences. All positions containing gaps and missing data were eliminated. There were 54 positions in the final dataset mainly corresponding to the DOF domain. Evolutionary analyses were conducted in MEGA7 (Kumar et al., 2016). DOFs from maritime pine have a red dot and their names are in bold. Abbreviations: At, *Arabidopsis thaliana*; Pp, *Pinus pinaster*; Os, *Oryza sativa*.

Figure S5. Phylogenetic tree of NAC transcription factor family in maritime pine and *Arabidopsis*. The protein sequences employed to infer the phylogenetic tree are presented in the Table S4. The protein sequence alignment was made with Muscle (Edgar, 2004). The evolutionary history was inferred using the Neighbor-Joining method (Saitou and Nei, 1987). The bootstrap consensus tree inferred from 1000 replicates is taken to represent the evolutionary history of the taxa analyzed (Felsenstein, 1985). Branches corresponding to partitions reproduced in less than 50% bootstrap replicates are collapsed. The evolutionary distances were computed using the Dayhoff matrix-based method (Schwarz and Dayhoff, 1979) and are in the units of the number of amino acid substitutions per site. The analysis involved 151 amino acid sequences. All positions with less than 80% site coverage were eliminated. That is, fewer than 20% alignment gaps, missing data, and ambiguous bases were allowed at any position. There were 261 positions in the final dataset. Evolutionary analyses were conducted in MEGA7 (Kumar et al., 2016). NACs from maritime pine have a red dot and their names are in bold.

Figure S6. Gene expression correlation network and functions. (**A**) Significant transcription factors for ammonium treatment that are hubs in a WGCNA gene correlation network and their connections with their correlated transcripts. The correlation cutoff value was |0,9|. Edges in red correspond to positive correlations. Edges in blue correspond to negative correlations. Red nodes are the hub TFs. Blue nodes are transcripts that always were significantly expressed in RDV. Some nodes were annotated with significant GO terms: turquoise nodes with GO:0022622 (Root system development); yellow nodes with GO:0010467 (Gene expression); orange nodes with GO:0042254 (Ribosome biogenesis); and pink nodes with GO:0010467 (Gene expression) and GO:0042254 (Ribosome biogenesis). (**B**) Functions of the correlated transcripts with hub TFs. Significant GO terms from Biological Processes category after a SEA analysis with a 0.4 cut-off value for dispensability.

Figure S7. Salicylic acid and jasmonic acid localization in root tip sections. The phytohormones were localized in root apex cuts of control and 3 mM NH_4_^+^ treated seedlings. The phytohormone identification and imaging was made using nano-particle assisted laser desorption/ionization mass spectrometry imaging (Nano-PALDI-MSI). (**A**) Salicylic acid (SA); (**B**) jasmonic acid (JA). Size bars correspond to 500 µm. Column graphs show relative hormone accumulations respect to control samples in the approximate LCM areas. Dashed lines mark hormone accumulation in the control roots.

Figure S8. Comparison between LCM and whole root transcriptomics data obtained from ammonium supply experiment with pine seedlings. DE transcripts were obtained from Ortigosa et al., 2021 (doi: 10.1101/2021.04.20.440618). (**A**) Expression of marker genes for whole root response to NH_4_^+^ supply in the tissues isolated through LCM. (**B**) Venn diagram with DE transcripts shared between LCM and whole root analyses. (**C**) LogFC data for shared DE transcripts between LCM and whole root approaches including the *antimicrobial peptide 1* (*PpAMP1*) and its identifiers at LCM dataset (pp_58007) and whole root (pp_58005).

Figure S9. Expression atlas of *PpDof1* in the tissues of one-month old seedlings (Cañas et al., 2017).

Figure S10. Expression atlas of *PpDof2* in the tissues of one-month old seedlings (Cañas et al., 2017).

Figure S11. Expression atlas of *PpDof11* in the tissues of one-month old seedlings (Cañas et al., 2017).

Figure S12. Expression atlas of *PpDof12* in the tissues of one-month old seedlings (Cañas et al., 2017).

**Dataset S1.** LCM low-input RNA-seq differential expression results. DE transcripts up-regulated are in highlighted red. DE transcripts down-regulated are in highlighted blue.

**Dataset S2.** LCM low-input RNA-seq clustering results of DE transcripts.

**Dataset S3.** LCM low-input RNA-seq functional enrichment of DE transcripts. Only conditions with significant functions are shown.

**Dataset S4.** Selected correlated transcripts with hub transcription factors for ammonium treatment.

**Dataset S5.** Functional enrichment of correlated transcripts with hub transcription factors for ammonium treatment.

## Notes

https://github.com/ceslobfer/Rootapp

### Competing Interest Statement

The authors have declared no competing interest.

