## Supplementary material for "Ammonium regulates the development of pine roots through hormonal crosstalk and differential expression of transcription factors in the apex": Table S1

**Table S1.** LCM low-input RNA sequencing results.

| **Sample** | **Raw bases (Gb)** | **Final bases (Gb)** | **Raw reads (M)** | **Final reads (M)** |
| --- | --- | --- | --- | --- |
| A1T24_3_RC | 11.92 | 9.9 | 79.47 | 68.01 |
| A1T24_3_RDC | 13.37 | 10.62 | 89.12 | 73.18 |
| A1T24_3_RDV | 11.64 | 9.6 | 77.58 | 66 |
| A1T24_3_RM | 12.83 | 10.58 | 85.52 | 72.75 |
| A1T24_C_RC | 13.39 | 10.99 | 89.24 | 75.87 |
| A1T24_C_RDC | 13.43 | 11.15 | 89.51 | 76.79 |
| A1T24_C_RDV | 12.41 | 10.3 | 82.73 | 70.78 |
| A1T24_C_RM | 11.66 | 9.69 | 77.76 | 66.58 |
| A3T24_3_RC | 12.16 | 9.96 | 81.08 | 68.83 |
| A3T24_3_RDC | 13.12 | 10.68 | 87.5 | 73.88 |
| A3T24_3_RDV | 13.23 | 10.83 | 88.2 | 74.94 |
| A3T24_3_RM | 12.82 | 10.67 | 85.46 | 73.55 |
| A3T24_C_RC | 13.22 | 10.86 | 88.13 | 75.16 |
| A3T24_C_RDC | 13.58 | 11.18 | 90.52 | 77.17 |
| A3T24_C_RDV | 12.48 | 10.38 | 83.23 | 71.75 |
| A3T24_C_RM | 12.9 | 10.62 | 85.98 | 73.28 |
| A4T24_3_RC | 13.71 | 11.44 | 91.42 | 78.82 |
| A4T24_3_RDC | 13.94 | 11.44 | 92.96 | 78.86 |
| A4T24_3_RDV | 14.87 | 12.19 | 99.11 | 84.48 |
| A4T24_3_RM | 12.45 | 10.32 | 82.97 | 71.08 |
| A4T24_C_RC | 15.67 | 12.79 | 104.46 | 88.36 |
| A4T24_C_RDC | 12.98 | 10.67 | 86.51 | 73.8 |
| A4T24_C_RDV | 12.47 | 10.39 | 83.13 | 71.54 |
| A4T24_C_RM | 12.89 | 10.47 | 85.4 | 71.87 |
