## Supplementary material for "Ammonium regulates the development of pine roots through hormonal crosstalk and differential expression of transcription factors in the apex": Table S2

**Table S2.** Primer list.

| **Primer Name** | **Identifier [*]** | **Sequence (5’ – 3’)** |
| --- | --- | --- |
| Amp-ATTB1.1 | - | TGCTCGGGGACAACTTTGTACAA |
| Amp-ATTB2.1 | - | GGCGGCCGCACAACTTTGTACAA |
| Amp-ATTB1.2 | - | TCGTCGGGGACAACTTT-GTACAAAAAAGTTGG |
| Amp-ATTB2.2 | - | GGCGGCCGCACAATTTGTACAA-GAAAGTTGGGTTTTTTT |
| polTdeg | - | AACAGTGGTATCAACGCAGAGTACT-TTTGTTTTTTTTTCTTTTTTTTTTVN |
| PugOligo–Adapter | - | AAGCAGTGGTATCAAC- GCAGAGTACGGGGG |
| qNADHGOGAT-F | pp_238923 | CATAACAAGCCACTCACATGCC |
| qNADHGOGAT-R | pp_238923 | CTTGGACCAGGTAGTTGATGCT |
| qppGS1b-F | pp_68481 | CCCAATTGTTTGTGGGGGATA |
| qppGS1b-R | pp_68481 | CTGAATGACAAACTAGACACTG |
| qppSAMS-F | pp_92107 | CTGTGCCCTCTTCATCCAGT |
| qppSAMS-R | pp_92107 | GCCATTACAGCCCACAGAAT |
| qppBLISTER-F | pp_170664 | TAAAGTTGTGTTTGCTGGGTTG |
| qppBLISTER-R | pp_170664 | TGCATGGTTTCAATGTTCTCTC |
| qppACO- F | pp_202153 | AATGCTTCATTTTCCACTGGTT |
| qppACO-R | pp_202153 | CGATGGAGCTCGATAAGAAAAT |
| qpp-AMP1-F | pp_58007 | AAAGGCGTTGCTCAGACCC |
| qpp-AMP1-R | pp_58007 | GCCAACTACTTAGACGTGCCT |
| qpp-SHR-F | pp_246622 | CAAGTGAAGTACCCGACCCC |
| qpp-SHR-R | pp_246622 | TTTGAAGCCCATCGACGTGA |
| qpp-NAC38-F | pp_72282 | TCTTATACTGCCTGCTTGCTTG |
| qpp-NAC38-R | pp_72282 | GTACGCATGGATTGGAGTGATA |
| qpp-NPF3.1-F | pp_58258 | TCCCCACCAACCTCAACAGAG |
| qpp-NPF3.1-R | pp_58258 | CCACAACGAAAGGCTTGTAGGT |
| qpp-CPI-F | pp_189071 | GATCCCCAAATGGATACAAAGA |
| qpp-CPI-R | pp_189071 | GTTTTCTTCCACATTGCACAAG |
| qpp-DOF12-F | pp_85381 | ACCCTTATGCCTGTAACAATGG |
| qpp-DOF12-R | pp_85381 | AGTACTGAACCCCGCAATACAT |
| qERCC-130-F | - | TCTGACGGGACAAGGGATCA |
| qERCC-130-R | - | ATTTCTGATATGGCGGCGGT |
| qERCC-2-F | - | GGGTCCATCAGTTGTCCGTA |
| qERCC-2-R | - | GTCCTTACAAGTCCGCTCCT |
| qERCC-96-F | - | CTTGCGCCAATTATCGAGCT |
| qERCC-96-R | - | AGTGTAGGACTCGTCGCATT |
| qERCC-4-F | - | ACATCTTCATAAGGGGTTGGGT |
| qERCC-4-R | - | TGGGGAAATTTGGGAAGCAGT |
| qERCC-46-F | - | TTCGGTGGCAGTATGGGATT |
| qERCC-46-R | - | ACAACACCAACGTCGCAAAAA |
| qERCC-136-F | - | CGCCAGTTTCCCGTGTATCT |
| qERCC-136-R | - | TCTTTCGGTCCAGTGCTTCC |
| qERCC-108-F | - | AGATATGCCATGCGTGCTGT |
| qERCC-108-R | - | CGCGTCGATAAGGTCACAGA |
| qERCC-116-F | - | GCTTATCGGGCCTGCTACAT |
| qERCC-116-R | - | GCTACCAGATCACCGCAGTT |
| qERCC-95-F | - | GGCTTAACCGCTATCGCTCT |
| qERCC-95-R | - | CGGCTTTGTGGGATGAGGTT |
| SlAP-F | pp_199988 | AGTATGCTAAGGAATCGTGCCT |
| SlAP -R | pp_199988 | GTCCATAATTACACACGAACAGA |
| SKP1/ASK1-F | unigene18128 | ATGCTGGACAGGCTTTGAAC |
| SKP1/ASK1-R | unigene18128 | GAGTTGCTCCGAGATCTTTACA |
