## Supplementary material for "Ammonium regulates the development of pine roots through hormonal crosstalk and differential expression of transcription factors in the apex": Methods S1

**Laser capture microdissection (LCM),** **RNA isolation and low-input RNA-seq**

LCM procedure was carried through as previously described (Cañas et al., 2014). One day before cryosectioning, root tip samples were tempered at -20°C. Thirty micrometers thick sections were cut using a cryostat (CM1950, Leica Biosysyems, Wetzlar, Germany) at -20°C and mounted on PET-membrane 1.4 µm steel frame (Leica Biosysyems, Wetzlar, Germany). The PET-membrane containing samples were stored at −80°C until use.

Before microdissection, samples were fixed in cold 70% ethanol for 30 seconds, embedding medium was removed in DEPC-treated water for 2 minutes and refixed in cold 100% ethanol for 2 minutes. Afterwards, samples were air dried and used for microdissection. A laser microdissector (LMD 7000, Leica, Wetzlar, Germany) was used for microdissection. The samples obtained were placed into 0.5 mL tube caps containing 20 μL of lysis buffer from the RNAqueous-Micro RNA Isolation Kit (Ambion, USA) and all samples were placed at −80°C. Four different tissue areas were isolated by microdissection approximatively corresponding to the root cap (RC), meristem (RM), developing cortex (RDC) and developing vessels (RDV) areas.

**cDNA amplification from LCM RNA samples**

One ng of total RNA was retrotranscribed and amplified to verify the expression analyses for several DE genes by RT-qPCR. The cDNA synthesis and amplification protocol was carried out using the Conifer RNA Amplification (CRA+) protocol previously described by Cañas *et al.* (2014). The amplification process was monitored using the ERCC RNA Spike-in kit (ThermoFisher, Waltham, MA, EEUU) according to manufacturer’s instructions. The primers used for cDNA synthesis and amplification are listed in the Table S2.

First‐strand cDNA synthesis 1 ng of total RNA was added to a mix containing 1 μM dNTPs and 1 μM polTdeg primer in a final volume of 7 μL. Reaction mixtures were incubated in a thermal cycler for 5 minutes at 65°C. Subsequently, temperature was decreased to 50°C keeping tubes in the thermal cycler at 50°C for 1-3 minutes. Afterwards, 3 μL RT mixture-1 (2 μL of 5× first‐strand buffer, 8.30 mM of MgCl2, 0.5 μL RevertAid H Minus Reverse Transcriptase (Thermo Scientific, Waltham, MA, EEUU) was added and reaction mix was gently mix by pipetting. All reaction tubes were incubated during 60 minutes at 50°C. When incubation time was ended, 10 μL of RT mixture-2 (2 μL of 5× first‐strand buffer, 5.1 μL nuclease‐free water, 8.30 mM MgCl_2_, 6 μM MnCl_2_ (100 mM), 1.95 mM PlugOligo‐Adapter primer and 0.5 μL RevertAid H Minus Reverse Transcriptase (Thermo Scientific, Waltham, MA, EEUU) was added to each tube. Reaction mixture was gently mixed by pipetting and incubated for 90 minutes at 42°C. The synthesized ss-cDNA was purified using the NucleoSpin® Gel and PCR Clean‐up kit (Macheray-Nagel, Düren, Germany) according to the manufacturer's instructions.

The ds-cDNA was obtained by nested PCR reactions using the primer pairs Amp-ATTB1.1 / Amp-ATTB2.1 and AmpATT-1.2 / Amp-ATTB2.2, respectively. First PCR (PCR-1) reactions were performed in a total volume of 25 μL, using 11.5 μL of purified ss-cDNA, 2X iProof™ HF Master Mix (BioRad, CA, USA) and 1 mM Amp-ATTB1.1 and Amp-ATTB2.1 primer pairs. The result obtained was diluted 35 times with nuclease-free water and several second PCR (PCR-2) reactions were performed using 1 μL of diluted ds-cDNA from PCR-1 step, 1 mM Amp-ATTB1.2 and Amp-ATTB2.2 primer pairs and 2X iProof™ HF Master Mix (BioRad, CA, USA) in a total volume of 50 μL. Several reactions were pooled and purified using NucleoSpin® Gel and PCR Clean‐up kit (Macheray‐Nagel, Düren, Germany) according to the manufacturer's instructions. The quantity of the amplified ds cDNA was determined using the Qubit 4 Fluorometer (Invitrogen, Paisley, UK) and Qubit dsDNA BR Assay Kit (Invitrogen, Paisley, UK) and the quality of the final amplifications were corroborated by 1% (p/v) agarose gel electrophoresis.
