## Supplementary figures and images for "Ammonium regulates the development of pine roots through hormonal crosstalk and differential expression of transcription factors in the apex"

### Figure S1

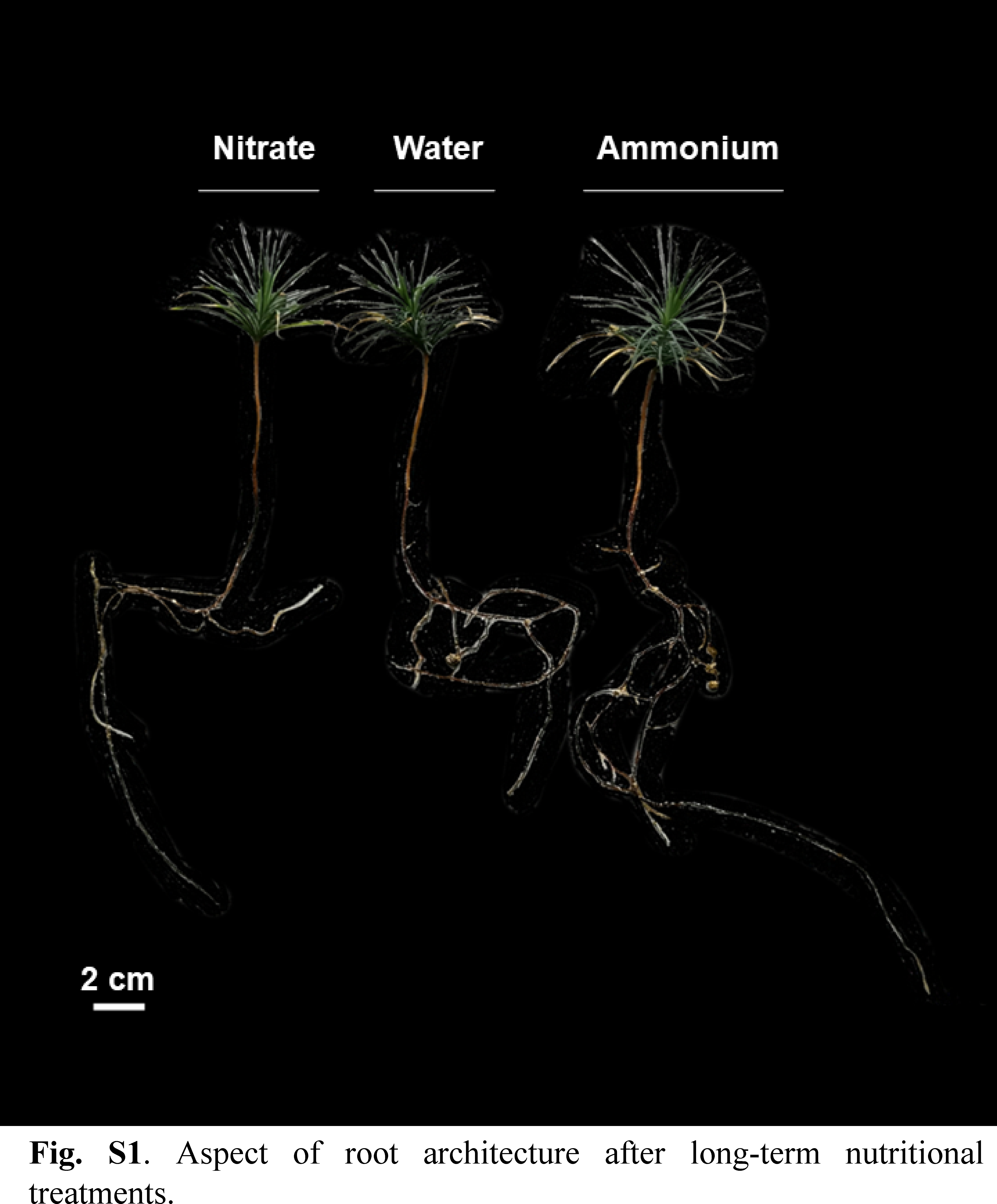

### Figure S2

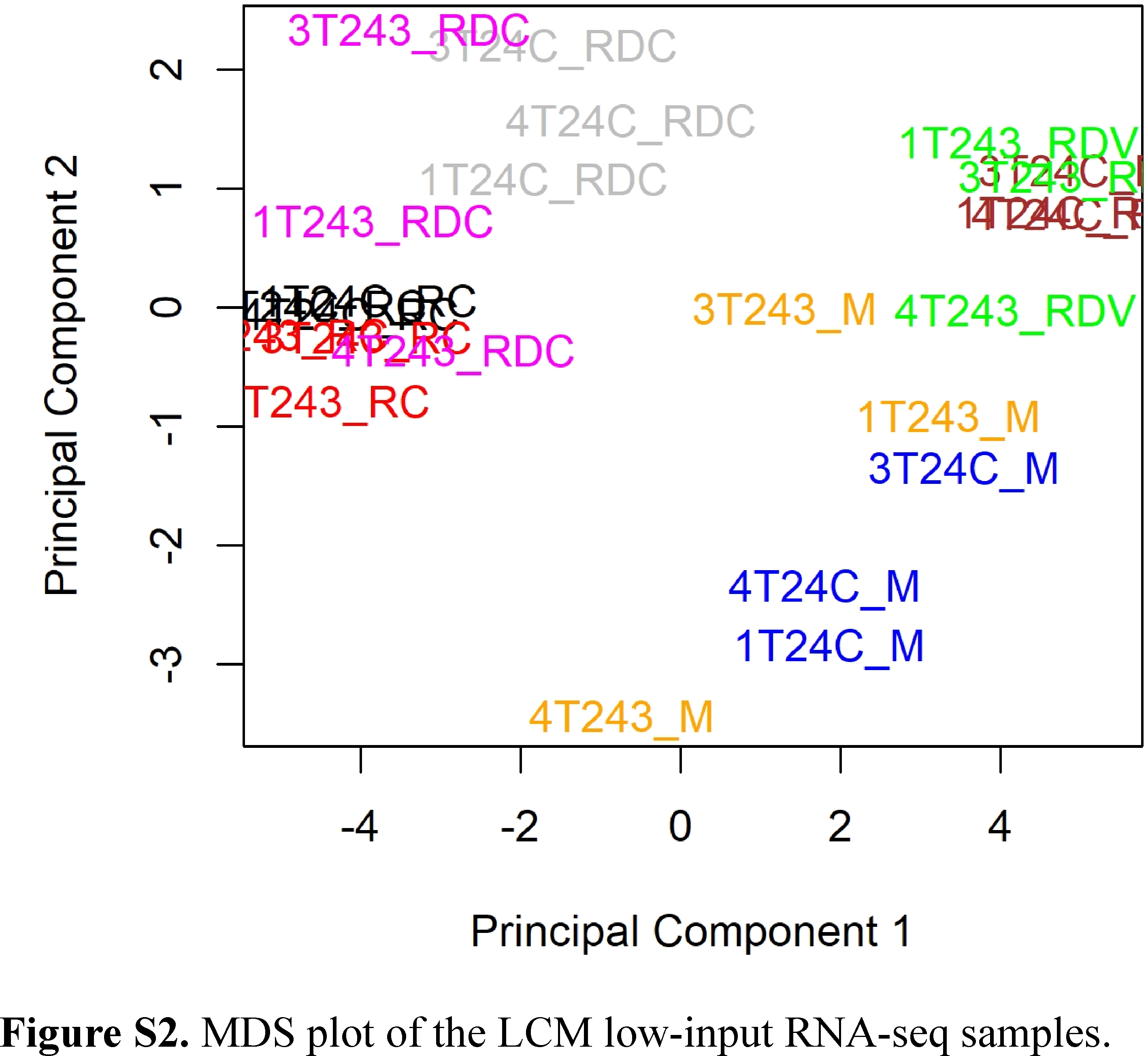

### Figure S4

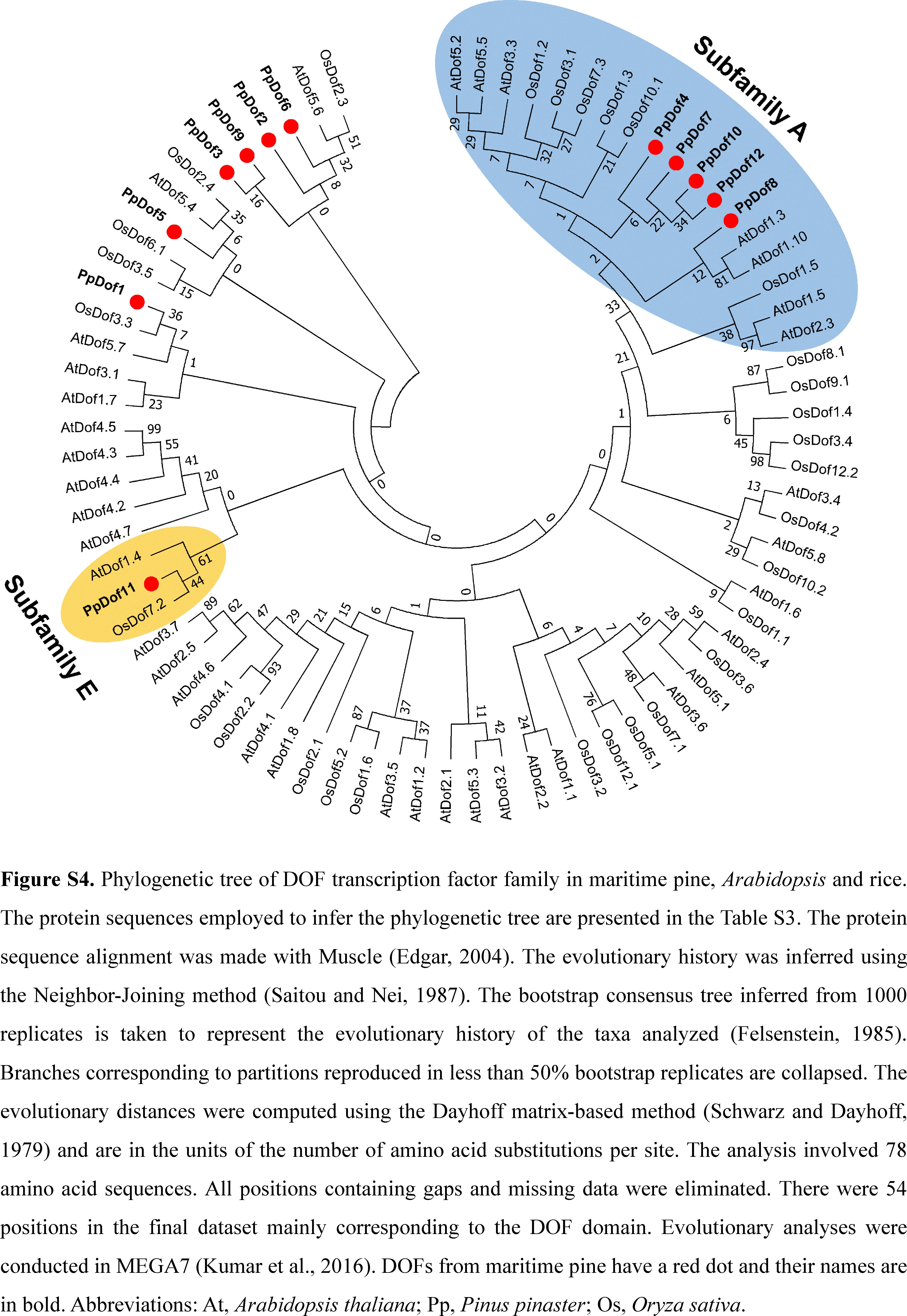

### Figure S5

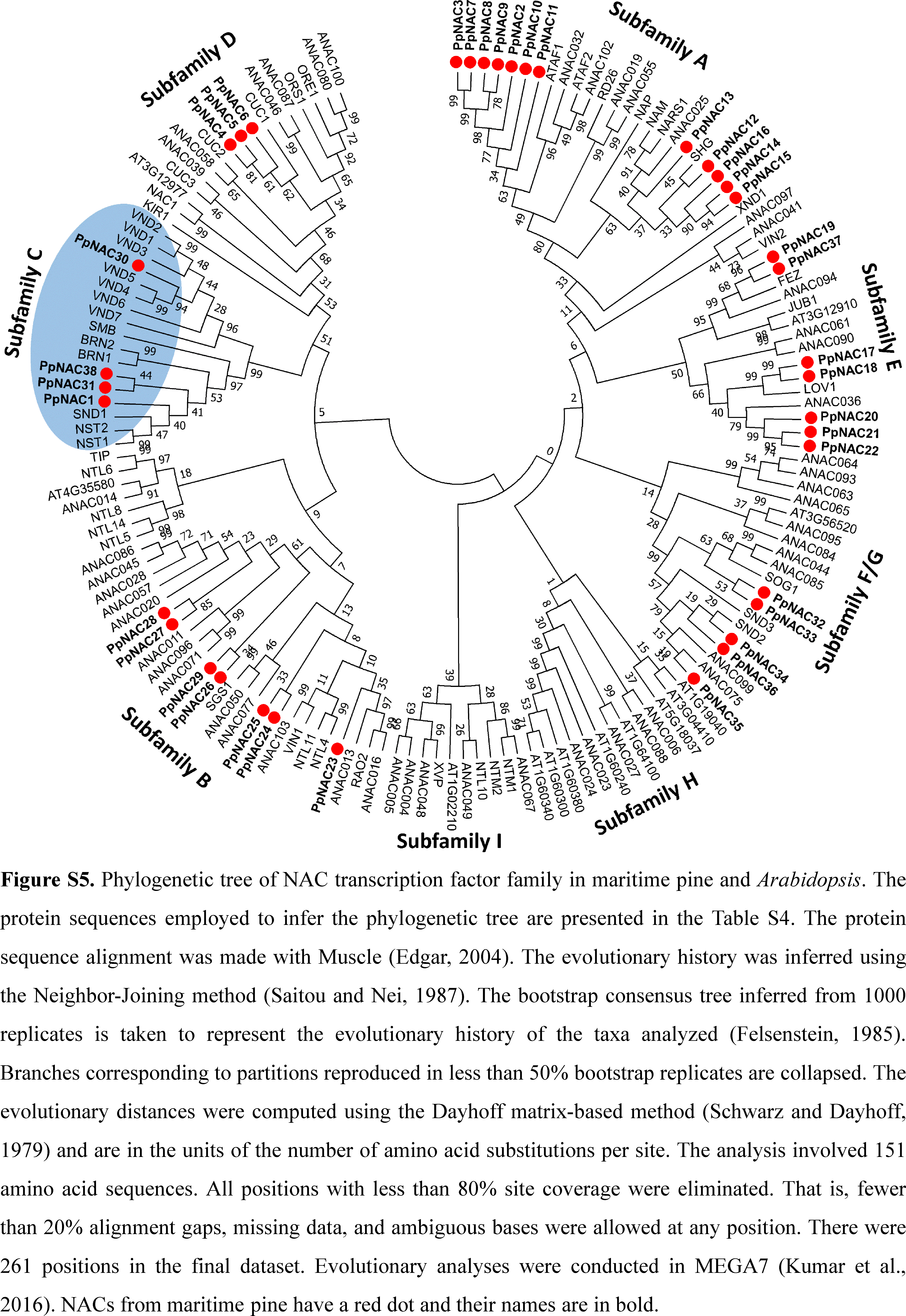

### Figure S7

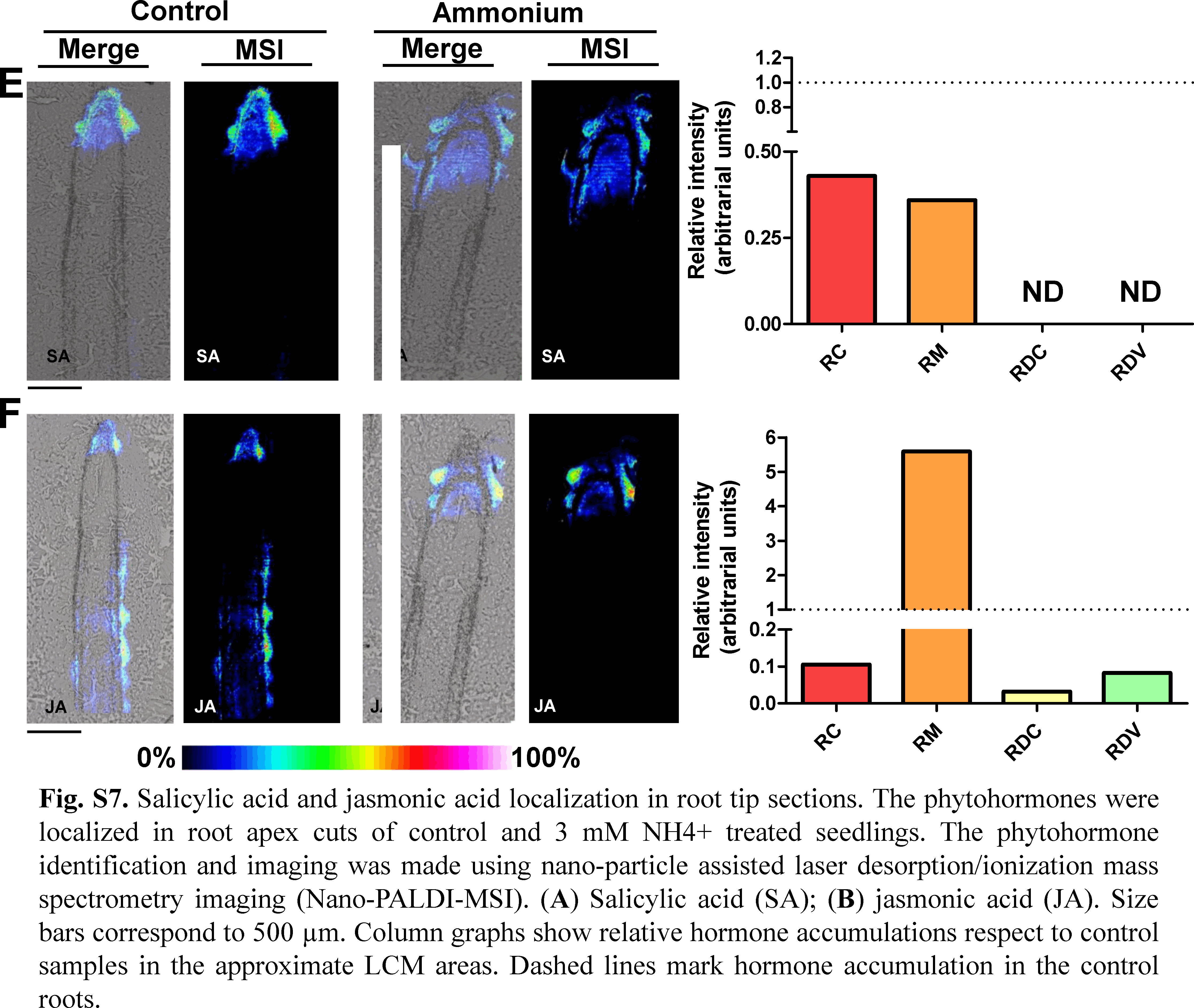

### Figure S8

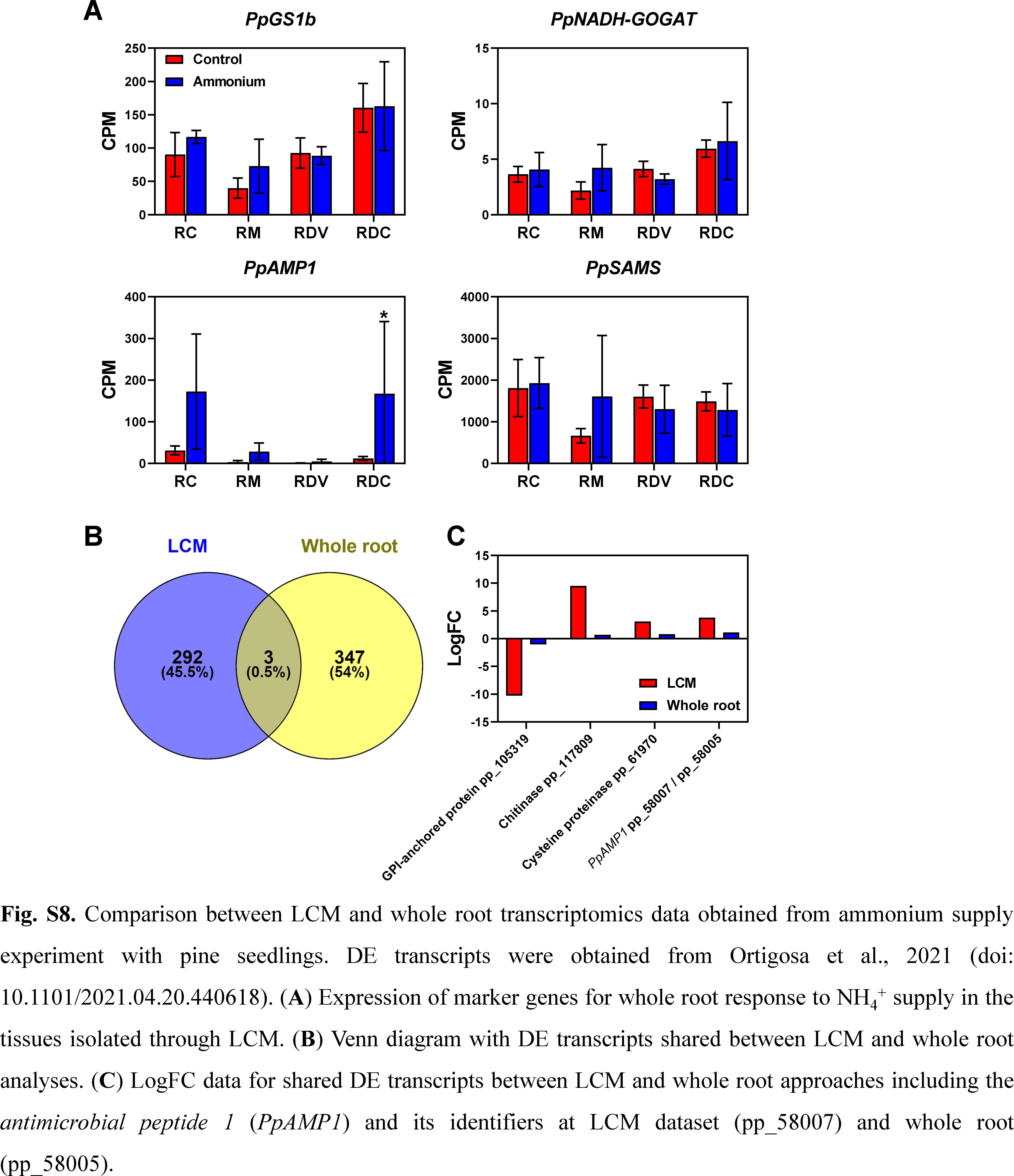

### Figure S9

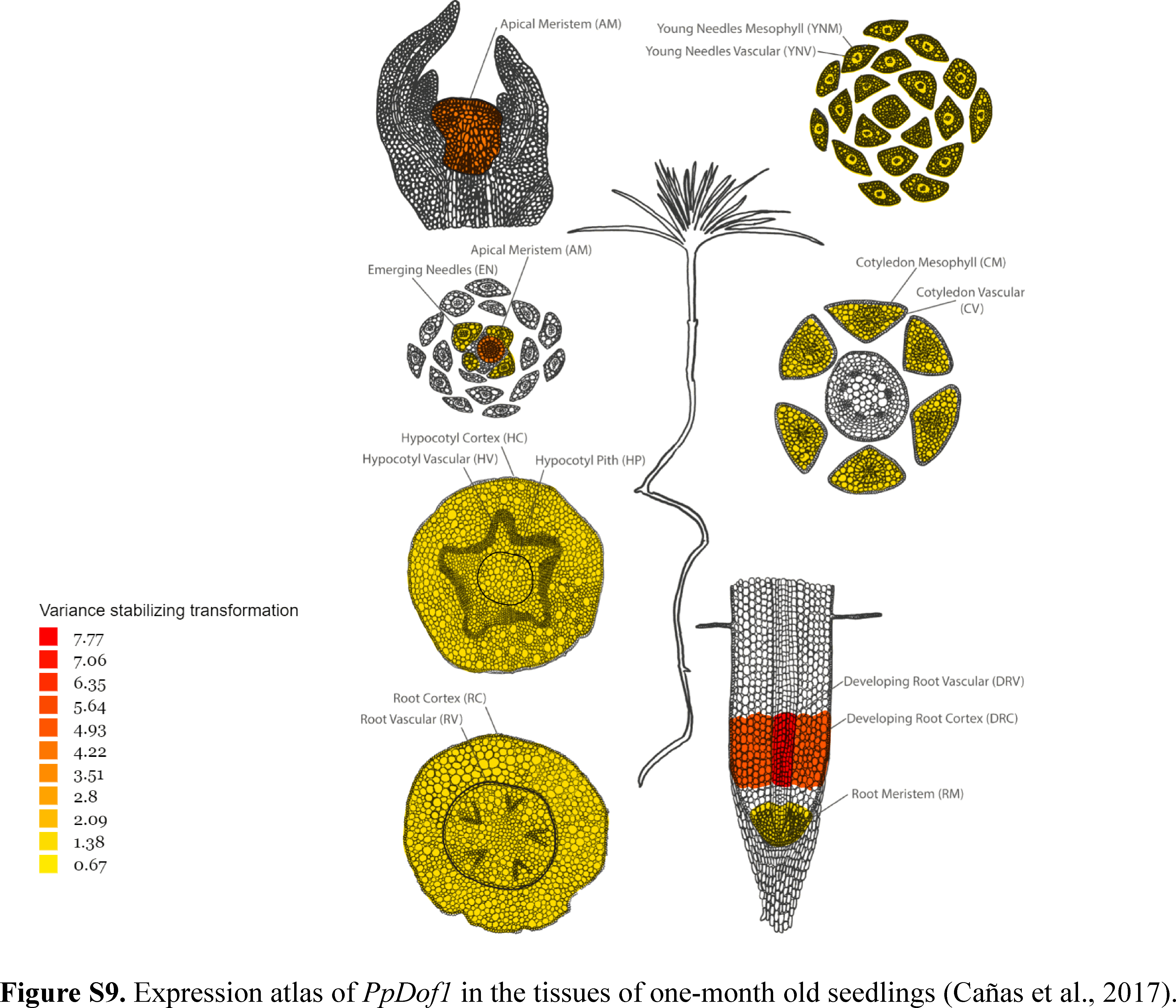

### Figure S10

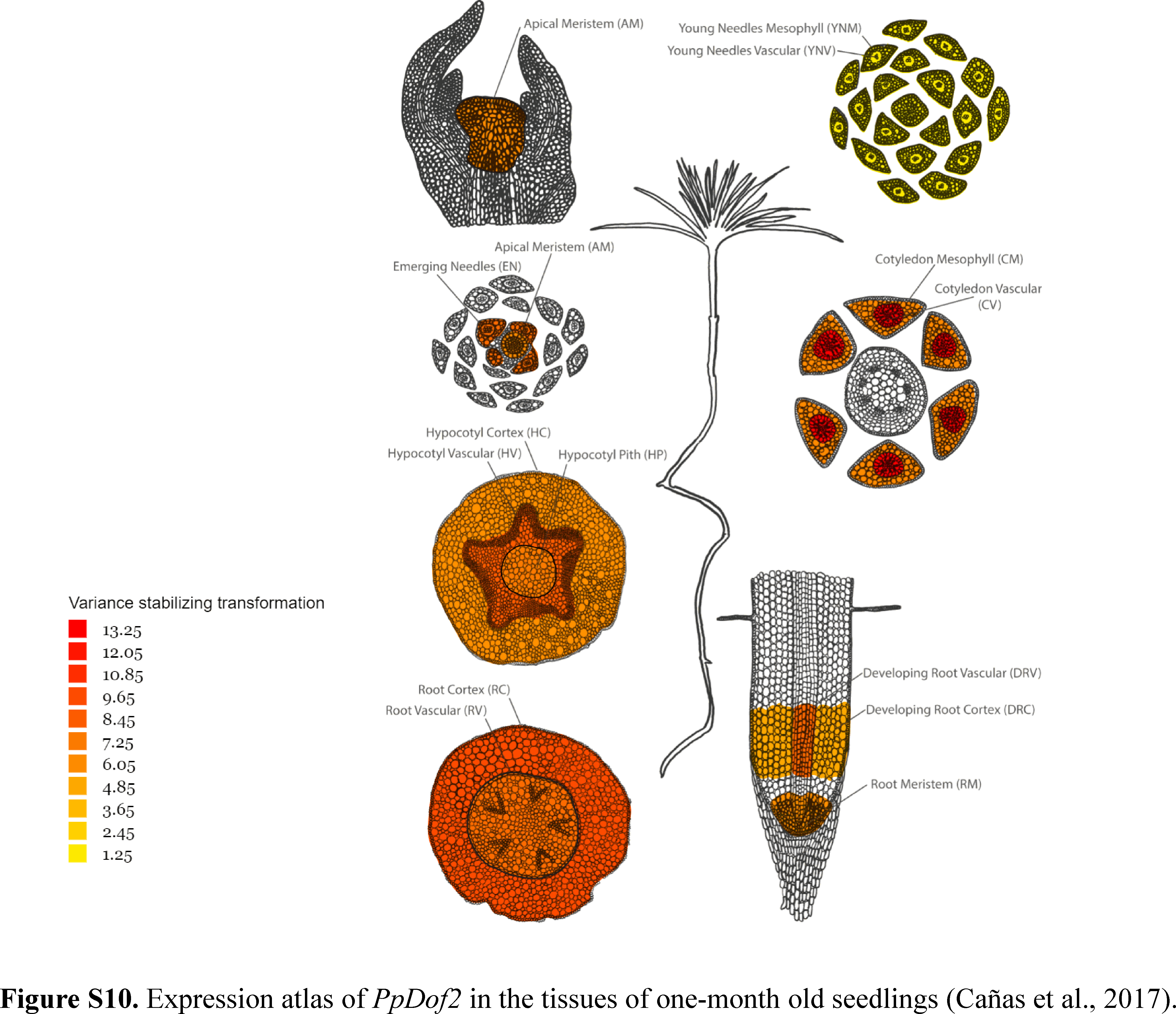

### Figure S11

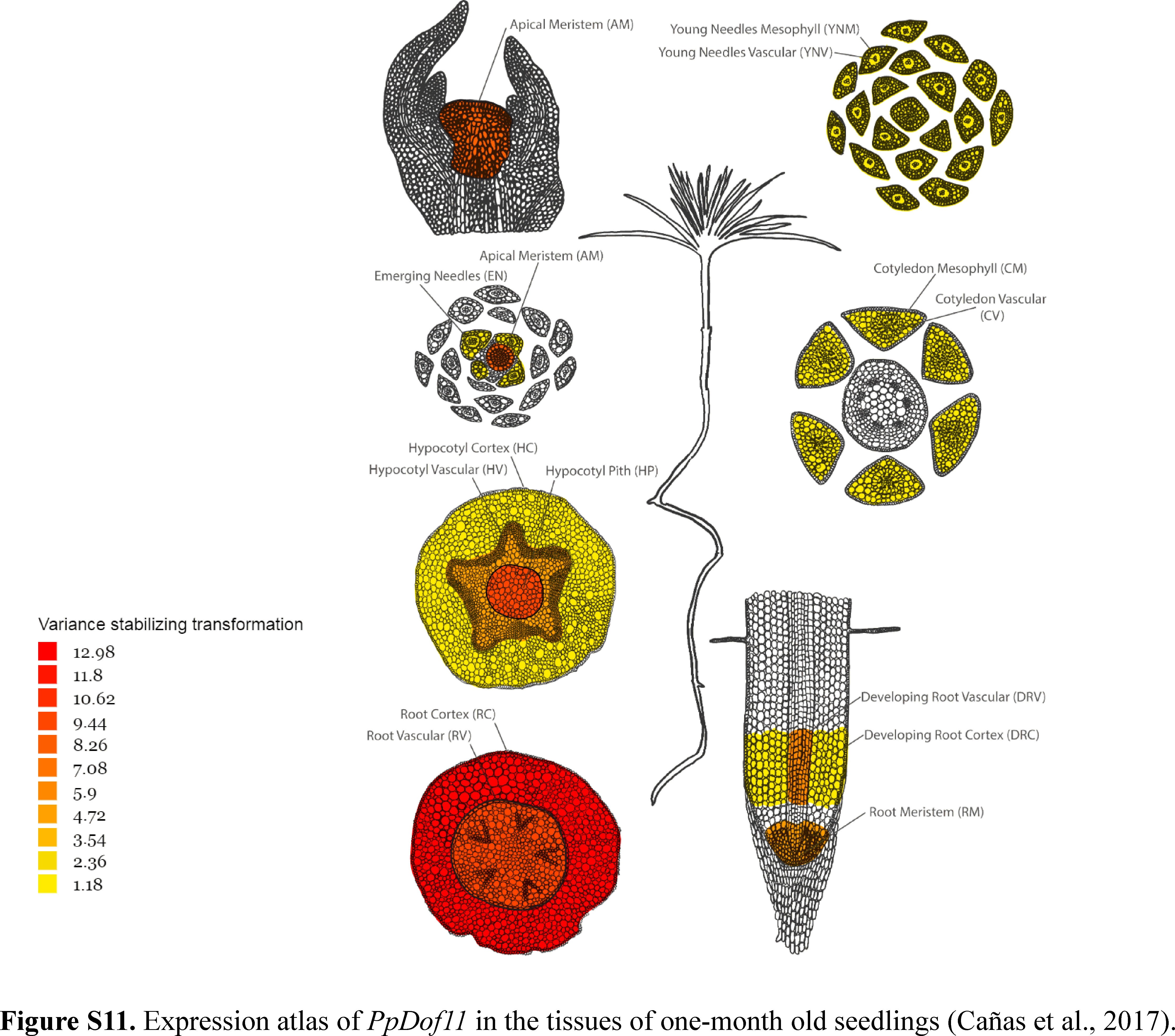

### Figure S12

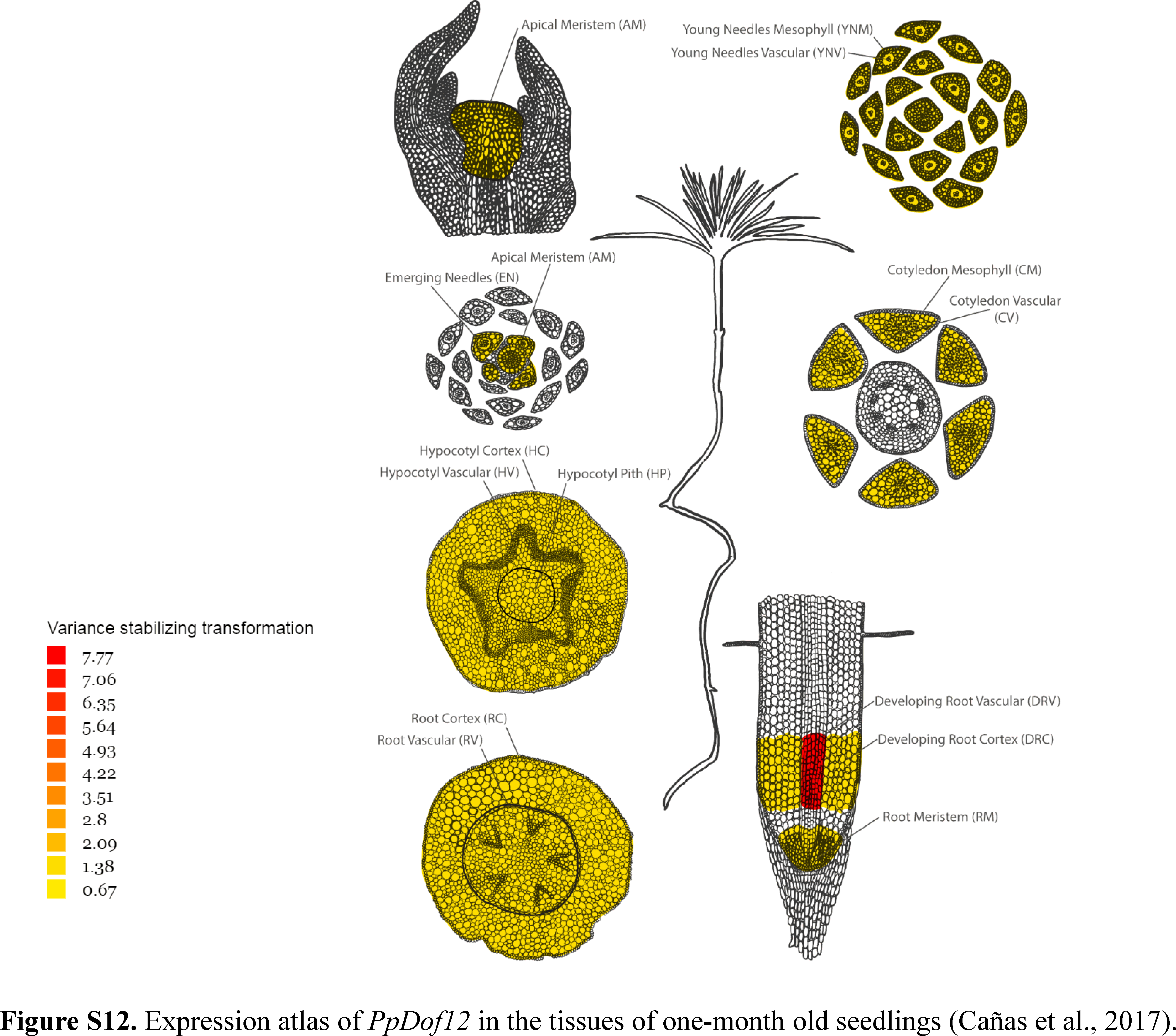
