## Supplementary material for "Ammonium regulates the development of pine roots through hormonal crosstalk and differential expression of transcription factors in the apex": Figure S3

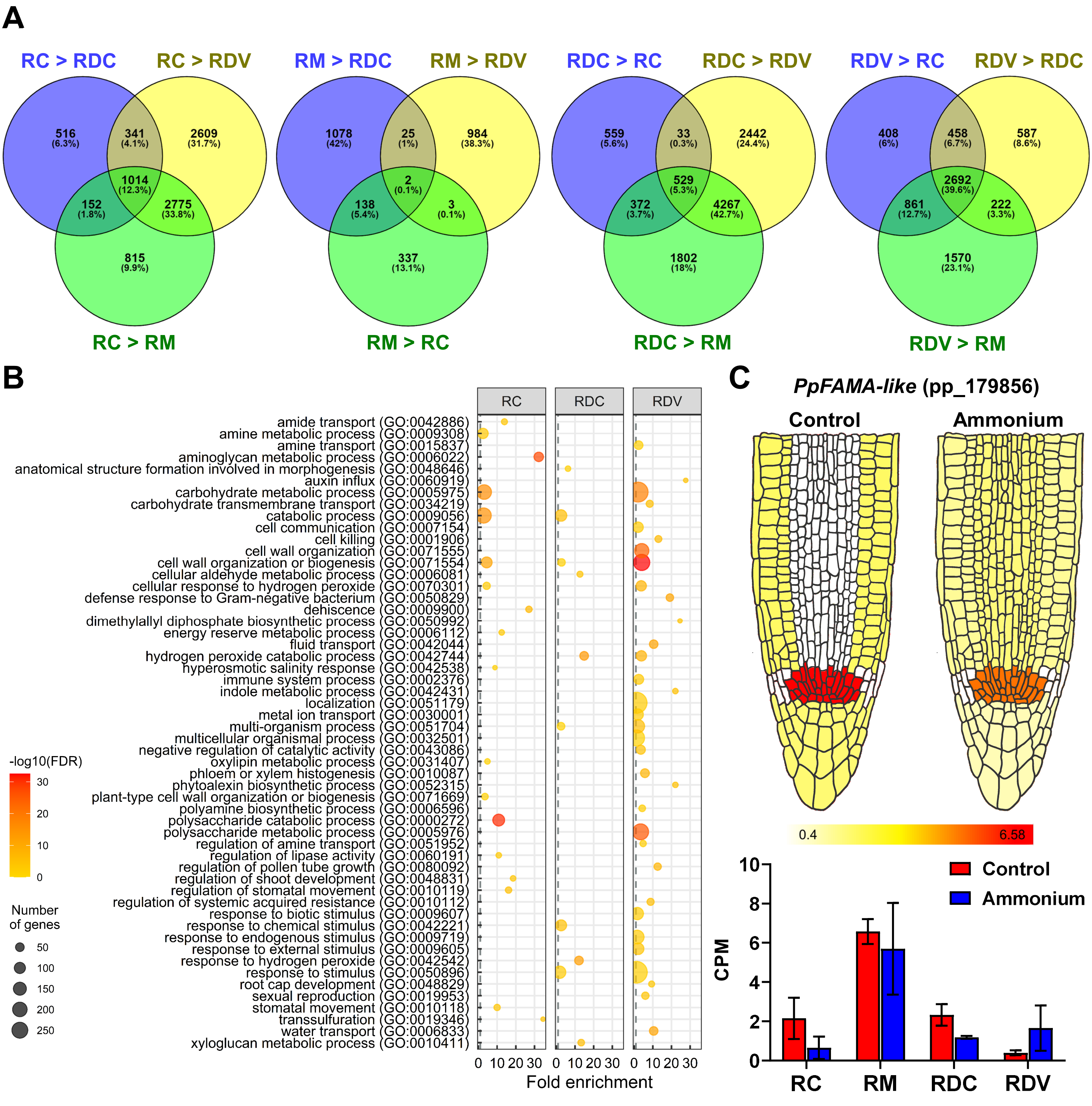


**Fig. S3.** RNA-seq differential expression analysis between tissues. (**A**) Venn diagrams shows the amounts of tissue-specific DE transcripts. (**B**) Functions of the transcripts that were always significantly expressed in a specific tissue. Significant GO terms from Biological Processes category after a SEA analysis with a 0.4 cut-off value for dispensability. (**C**) Expression of transcription factor *PpFAMA-like* (pp_179856) in the different isolated tissues of the root apex. *PpFAMA-like* was differentially expressed in the RM respect to the rest of tissues but not in the nutritional treatment comparison.
