## Supplementary material for "Ammonium regulates the development of pine roots through hormonal crosstalk and differential expression of transcription factors in the apex": Figure S6

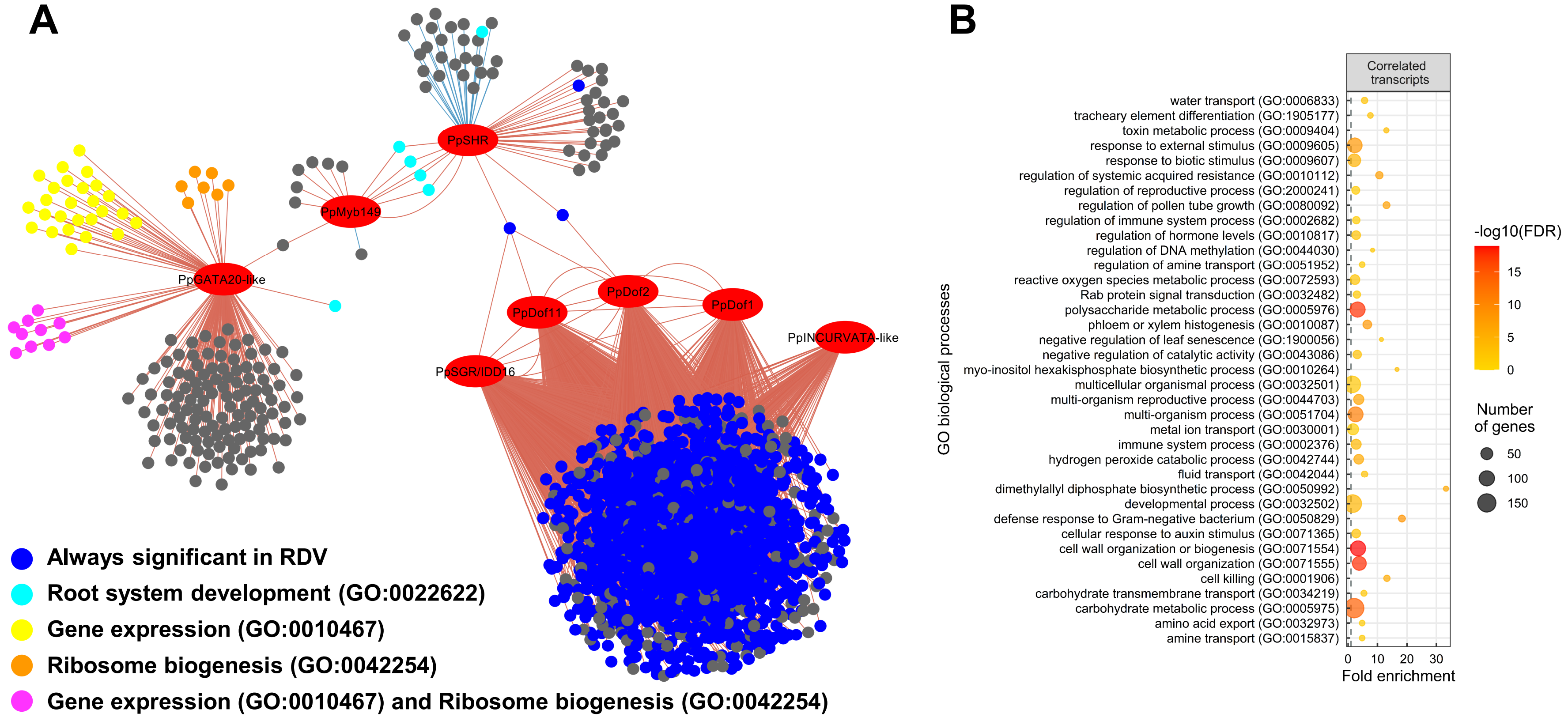


**Fig. S6.** Gene expression correlation network and functions. (**A**) Significant transcription factors for ammonium treatment that are hubs in a WGCNA gene correlation network and their connections with their correlated transcripts. The correlation cutoff value was |0,9|. Edges in red correspond to positive correlations. Edges in blue correspond to negative correlations. Red nodes are the hub TFs. Blue nodes are transcripts that always were significantly expressed in RDV. Some nodes were annotated with significant GO terms: turquoise nodes with GO:0022622 (Root system development); yellow nodes with GO:0010467 (Gene expression); orange nodes with GO:0042254 (Ribosome biogenesis); and pink nodes with GO:0010467 (Gene expression) and GO:0042254 (Ribosome biogenesis). (**B**) Functions of the correlated transcripts with hub TFs. Significant GO terms from Biological Processes category after a SEA analysis with a 0.4 cut-off value for dispensability.
